# CDK8 remodels the tumor microenvironment to resist the therapeutic efficacy of targeted KRAS^G12D^ inhibition in pancreatic ductal adenocarcinoma

**DOI:** 10.1101/2025.01.29.635543

**Authors:** Kathleen M. McAndrews, Krishnan K. Mahadevan, Bingrui Li, Amari M. Sockwell, Sami J. Morse, Patience J. Kelly, Sarah I. Patel, Michelle L. Kirtley, Barbara A. Moreno Diaz, Hengyu Lyu, David H. Peng, Xunian Zhou, Hikaru Sugimoto, Aaron A. Bickert, Lakshmi Kavitha Sthanam, Meagan R. Conner, Shreyasee V. Kumbhar, Kent A. Arian, Yasaman Barekatain, Francesca Paradiso, Paola A. Guerrero, Vincent Bernard, Navid Sobhani, Alondra N. Camacho-Acevedo, Kiara E. Bornes, Phuong Thao Tran, Anirban Maitra, Timothy P. Heffernan, Raghu Kalluri

**Author notes:** Corresponding author: Raghu Kalluri, MD, PhD.

## Abstract

Mutations in *KRAS* are a dominant driver of pancreatic ductal adenocarcinoma (PDAC), with over 40% of PDAC patients presenting with *KRAS^G12D^* mutations. The recent development of small molecule inhibitors targeting KRAS^G12D^ has enabled targeting of mutant KRAS signaling and suppression of PDAC; however, the contribution of the tumor microenvironment (TME) to the sustained therapeutic efficacy of KRAS^G12D^ inhibition and mechanism/s of resistance to KRAS^G12D^ suppression remain to be elucidated. Here, we employed spatial transcriptomics, single cell RNA sequencing, and CODEX-based spatial proteomics to evaluate cancer cell intrinsic and extrinsic responses to KRAS^G12D^ inhibition with MRTX1133. While KRAS^G12D^ inhibition initially increases CD11c^+^ cells with impactful T cell infiltration within proximity to cancer cells, long-term treatment with MRTX1133 resulted in reversal of the immune responses leading to KRAS^G12D^ therapy resistance promoted by CDK8, a multiprotein mediator complex associated kinase. CDK8 imparts resistance in part through induction of downstream CXCL2 chemokine secretion, inhibition of FAS expression, and remodeling of the TME to promote immune evasion. Targeting CDK8 by itself and in combination with αCTLA-4 immunotherapy overcomes resistance to KRAS^G12D^ inhibition with prolonged survival with translational implications.

## Introduction

Pancreatic ductal adenocarcinoma is characterized by tissue-specific genetic events, including frequent mutations in *KRAS*. Specifically, *KRAS^G12D^* mutations occur in approximately 40% of patients^1^ and genetically engineered mouse models (GEMMs) have revealed that oncogenic KRAS (KRAS*) is critical for PDAC initiation and maintenance,^2–6^ suggesting that targeting KRAS* may provide therapeutic benefits. KRAS* promotes the expression of pERK and reprograms cancer cell metabolism to promote glucose uptake and glycolysis, which is thought to be critical for supporting epithelial proliferation.^4,6^ In addition to its role in promoting cancer cell growth and other cancer cell intrinsic signaling pathways, KRAS* has also been implicated in remodeling of the tumor microenvironment (TME). KRAS* induces desmoplasia in part through the activation of the Hedgehog signaling pathway in epithelial cells^6^ and by promoting the expression of inflammatory genes.^7^ KRAS* promotes the epigenetic silencing of FAS, preventing CD8^+^ T cell-mediated induction of cancer cell apoptosis,^3,8^ suggesting that KRAS* generates an immunosuppressive microenvironment. The recent development of KRAS* small molecule inhibitors has enabled therapeutic targeting of KRAS*, and such inhibitors impact cancer cell intrinsic signaling as well as reprogram the TME.^9,10^ MRTX1133 was developed as a small molecule non-covalent inhibitor of KRAS^G12D^ that displays selective binding to the GDP-bound inactive form of KRAS^G12D^.^11^ MRTX1133 has been shown to inhibit proliferation and downstream KRAS signaling, ascertained by measuring the surrogate readouts of KRAS activity pERK and pS6, in several KRAS^G12D^ mutant PDAC cell lines and animal models^9–12^. In addition to the impact of MRTX1133 on cancer cell autonomous signaling and proliferation, previous studies have implicated a role for the immune TME in facilitating responses to MRTX1133 and provided a rationale for combining MRTX1133 with immune checkpoint blockade (ICB).^9,10,12^

While KRAS* small molecule inhibition has demonstrated effective tumor control in xenograft, syngeneic orthotopic models and spontaneous mouse models, prolonged KRAS* inhibition ultimately leads to tumor relapse,^9–11^ similar to what has been reported with genetic KRAS* suppression and inhibition of the KRAS* and receptor tyrosine kinase (RTK) signaling pathways with small molecule inhibitors.^13–15^ Earlier studies suggested that PDAC cancer cells that are resistant to genetic KRAS* suppression are metabolically dependent on oxidative phosphorylation and in some contexts display amplification of *Yap*, which acts to promote KRAS*-independent growth.^14,15^ Moreover, *Hdac5* was identified as promoting KRAS*-independent growth through increased CCL2 in cancer cells, which acts to promote the infiltration of CCR2^+^ macrophages.^16^ Several potential mechanisms of KRAS* small molecule inhibitor resistance have been proposed in the context of resistance to small molecule inhibition of KRAS^G12C^, including mutations in the drug binding pocket of kinases, increased signaling through alternative pathways to promote growth, and dysregulation of the cell cycle;^17–19^ however, there is emerging evidence that subtypes of *KRAS* mutations may have distinct functional outcomes. KRAS* allele specific progression of PanIN lesions occurs in GEMMs, suggesting that common and divergent mechanisms of KRAS* inhibition may exist.^20^ Moreover, KRAS^G12C^ and KRAS^G12D^ have different intrinsic and GAP-stimulated GTP hydrolysis rates,^21^ which may impact small molecule inhibitor binding to the inactive and active states of KRAS* and overall KRAS* inhibition efficacy.

Mechanism(s) of resistance to small molecule inhibition of KRAS^G12D^ remain to be unraveled. Here, we identify CDK8 as promoting KRAS* inhibition resistance in PDAC through its suppression of FAS expression and induction of CXCL2, which act to promote immunosuppression. Combined inhibition of KRAS* with CDK8 or the CXCL2 receptor CXCR2 inhibits tumor growth and prolongs survival, indicating a role for the immune microenvironment in shaping resistance to a small molecule inhibitor of KRAS*.

## Results

### Long term KRAS* inhibition leads to therapy resistance

To evaluate the impact of long-term KRAS* inhibition on PDAC progression and resistance, multiple mouse models of PDAC were treated with MRTX1133. *Trp53^F/+^; LSL-Kras^G12D/+^; p48-Cre* (PKC-HY19636) cells were orthotopically implanted in the pancreata of syngeneic C57BL/6 mice and mice treated with the non-covalent small molecule KRAS* inhibitor MRTX1133 (**Fig. 1A**). Mice treated with MRTX1133 (30 mg/kg BID i.p.) demonstrated significantly reduced tumor burden at one-week post-treatment initiation (MRTX1133 sensitive); however, mice with continued MRTX1133 treatment (MRTX1133 resistant, approximately 2-3 weeks of treatment) ultimately relapsed and displayed similar tumor burden to control (vehicle) treated mice (**Fig. 1B**). Histological assessment further confirmed tumor relapse in MRTX1133 resistant mice (**Fig. 1C-D**). Reduced pERK^+^ cells were observed in MRTX1133 treated tumors at early timepoints, indicative of suppression of KRAS signaling, whereas pERK^+^ cells in MRTX1133 resistant tumors were heterogeneous, with some tumors having high numbers and others having lower numbers of pERK^+^ cells (**Fig. 1E-F**). Costaining for CK19 and pERK revealed specific downregulation of pERK in CK19^+^ cancer cells in MRTX1133 sensitive tumors and heterogeneous responses in MRTX1133 resistant tumors (**Supplemental Fig. 1A-B**), similar to IHC results (**Fig. 1E-F**). Moreover, CK19^−^pERK^+^ cells were not changed in MRTX1133 treated mice (**Supplemental Fig. 1A-B**), supporting specific targeting of mutant KRAS^G12D^ as previously reported^9,11^. The *Kras^G12D^* mutation was detected in vehicle, MRTX1133 sensitive, and MRTX1133 resistant tumors (**Fig. 1G**), supporting that resistance is associated with outgrowth of *Kras^G12D^* mutant cells.

**Figure 1:**
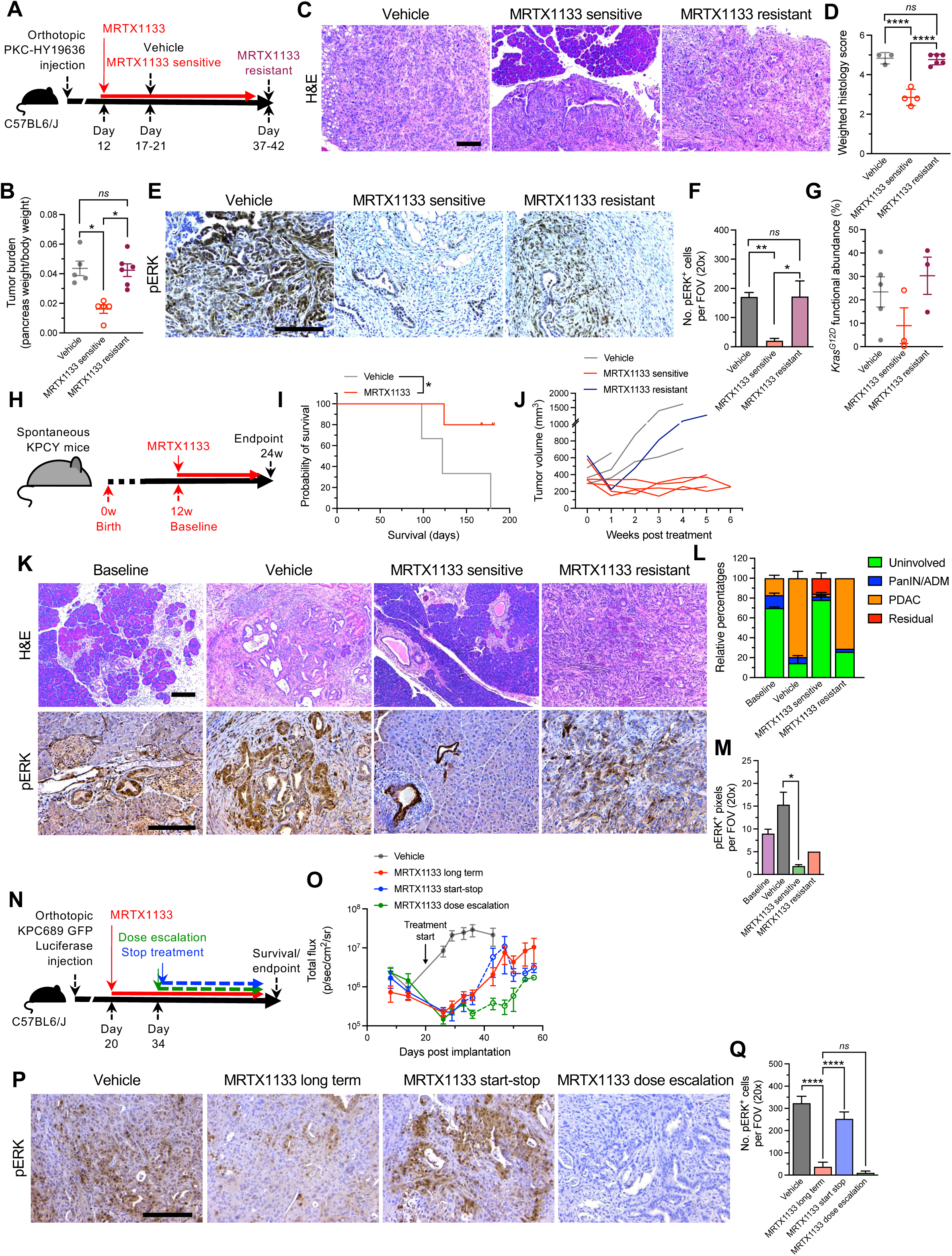
Murine PDAC models demonstrate resistance to KRAS^G12D^ inhibition. (**A**) Schematic of orthotopic PKC-HY19636 tumors treated with MRTX1133. (**B**) Tumor burden of PKC-HY19636 tumors. Vehicle, n=5; MRTX1133 sensitive, n=5; MRTX1133 resistant, n=6. (**C-D**) Representative H&E images (**C**) and quantification (**D**) of orthotopic PKC-HY19636 tumors. Vehicle, n=3; MRTX1133 sensitive, n=4; MRTX1133 resistant, n=6. (**E-F**) Representative pERK staining (**E**) and quantification (**F**) of orthotopic PKC-HY19636 tumors. Vehicle, n=5; MRTX1133 sensitive, n=6; MRTX1133 resistant, n=6. (**G**) ddPCR for *Kras^G12D^* functional abundance in PKC-HY19636 orthotopic tumors. Vehicle, n=5; MRTX1133 sensitive, n=3; MRTX1133 resistant, n=3. (**H**) Schematic of spontaneous KPCY GEMM tumors treated with MRTX1133. (**I**) Survival curve of KPCY GEMM mice (days post-birth). Vehicle, n=3; MRTX1133, n=5. (**J**) Tumor volume of KPCY GEMMs measured by MRI. Vehicle, n=3; MRTX1133 sensitive, n=4; MRTX1133 resistant, n=1. (**K-M**) Representative H&E and pERK images (**K**), histological assessment (**L**), and pERK quantification (**M**). Baseline, n=2; Vehicle, n=3; MRTX1133 sensitive, n=4; MRTX1133 resistant, n=1. Scale bars, 100 μm. (**N**) Schematic of orthotopic KPC689 tumors treated with MRTX1133. (**O**) Bioluminescence of orthotopic KPC689 tumors in C57BL6/J mice over time. Treatment was initiated at day 20 (Vehicle: n=8, MRTX1133 long term: n=9, MRTX1133 start-stop: n=8, MRTX1133 dose escalation: n=9). Dashed lines denote where treatment was stopped for MRTX1133 start-stop and dosage increased for MRTX1133 dose escalation. (**P-Q**) Representative pERK images (**P**) quantification (**Q**) of orthotopic KPC689 tumors. Vehicle: n=4, MRTX1133 long term: n=5, MRTX1133 start-stop: n=3, MRTX1133 dose escalation: n=6. Kruskall-Wallis with Dunn’s multiple comparisons test performed in (B) and (F). One-way ANOVA with Tukey’s multiple comparisons test performed in (D). Log-rank test performed in (I). Unpaired t-test with Welch’s correction performed in (M). One-way ANOVA with Dunnett’s multiple comparisons test performed in (Q). Scale bars, 100 μm. Data are presented as mean ± s.e.m. * P < 0.05, *** P < 0.001, **** P < 0.0001, *ns*: not significant.

In the context of spontaneously occurring *LSL-Trp53^R172H/+^*; *LSL-Kras^G12D/+^; Pdx1-Cre; LSL-EYFP* (KPCY) tumors, increased survival and tumor growth control was observed in response to MRTX1133, albeit in a limited number of mice (30 mg/kg BID i.p., **Fig. 1H-I**). Like orthotopic models, tumor regrowth following MRTX1133 treatment was observed at 2-5 weeks post-treatment initiation (MRTX1133 resistant, **Fig. 1J**, **Supplemental Fig. 1C**) without impacting bodyweight during treatment (**Supplemental Fig. 1D**). Tumors that responded to MRTX1133 treatment (MRTX1133 sensitive) had increased uninvolved tissue relative to vehicle treated tumors as evaluated by H&E staining (**Fig. 1K-L**). EYFP, a lineage marker of acinar cells and cancer cells, immunostaining was performed and evaluated in conjunction with H&E stains to identify the presence of cancer cells derived from the *Pdx1*^+^ (acinar) cell lineage (**Supplemental Fig. 1E**). pERK was suppressed in MRTX1133 sensitive tumors with only a marginal upregulation in the resistant tumor (**Fig. 1K, 1M**), suggesting suppression of KRAS^G12D^ in spontaneous PDAC models.

A second orthotopic model was employed to confirm emergence of resistance to KRAS* inhibition, wherein KPC689 GFP (green fluorescent protein)-luciferase orthotopic tumor bearing mice were treated with MRTX1133 and tumor kinetics were monitored with IVIS imaging (**Fig. 1N**). MRTX1133 treatment was associated with initial tumor regression followed by relapse within 14 days in the KPC689 GFP-luciferase model (**Fig. 1O, Supplemental Fig. 1F-G**). After 14 days of treatment, one group of mice was maintained on MRTX1133 treatment (MRTX1133 long term), one group of mice were enrolled in a higher dose of MRTX1133 (MRTX1133 dose escalation, 60 mg/kg BID i.p.) and treatment stopped on another group of mice (MRTX1133 start-stop, **Fig. 1N**). Bodyweight was similar across treatment groups following treatment enrollment (**Supplemental Figure 1H**). Dose escalation initially controlled tumor growth but ultimately was associated with resistance to KRAS* inhibition and PDAC regrowth (**Fig. 1N-O, Supplemental Fig. 1F-G, Supplemental Fig. 2A-B**). Dose escalation was also associated with non-PDAC related death, suggesting that toxicity may exist at higher doses of MRTX1133 despite no overt changes in bodyweight (**Supplemental Fig. 1H**). Treatment cessation had no impact on tumor growth (**Fig. 1O, Supplemental Fig. 1F-G, Supplemental Fig. 2A-B**). MRTX1133 long term and MRTX1133 dose escalation tumors both demonstrated reduced pERK^+^ cells, whereas pERK^+^ cells were increased in vehicle and MRTX1133 start and stop treatment tumors (**Fig. 1P-Q**). Emergence of resistance was further validated in another PDAC mouse model using the KPC-T cell line. This cell line was established after serial transplantation of a tumor from KPC mice (KPC-T)^22^. KPC-T cells were orthotopically implanted into C57BL/6 mice and treated with MRTX1133 (**Supplemental Fig. 2C**). MRTX1133 significantly increased the overall survival of mice and led to control of tumor growth at 1 week following treatment; however, prolonged treatment was associated with tumor reemergence (**Supplemental Fig. 2D-F**). MRTX1133 sensitive tumors, 1 week following MRTX1133 treatment, showed reduced tumor burden by histological analysis and pERK suppression, whereas increased tumor burden and sustained suppression of pERK was observed in MRTX1133 resistant tumors (**Supplemental Fig. 2E-G**), supporting the emergence of resistance to MRTX1133 in multiple models of PDAC.

### KRAS* inhibitor resistant cancer cells upregulate CDK8

To evaluate the transcriptional changes resulting from MRTX1133 treatment, single cell RNA sequencing (scRNA-seq) was employed, and ductal/cancer cells specifically analyzed with 6 transcriptional clusters of cells identified (**Fig. 2A-B, Supplemental Fig. 3A-C**). InferCNV was performed to identify cancer cells within the ductal/cancer cell cluster, with tumor-derived B cells and T and NK cells used as a reference (**Supplemental Fig. 4A**). Cluster 6 specifically lacked overt deletions and amplifications, like the reference cells, suggesting that this cluster likely consists of untransformed ductal cells (**Supplemental Fig. 4A**). In MRTX1133 sensitive, cluster 6 is the primary cluster represented, supporting effective elimination of cancer cells with treatment (**Fig. 2B**). Gene set enrichment analysis (GSEA) of MRTX1133 resistant tumors compared to vehicle tumors revealed downregulation of a number of pathways previously associated with KRAS*, including downregulation of hypoxia, glycolysis, KRAS signaling, and mTORC1 signaling, and upregulation of MYC targets and oxidative phosphorylation (**Fig. 2C, Supplemental Fig. 4B**).^4,15,23^ Further analysis of KRAS pathway related genes, *Dusp6* and *Spry4* revealed downregulation with MRTX1133 treatment that was maintained in resistant tumors (**Fig. 2D**), indicating effective suppression of KRAS* signaling. Previous studies have implicated a rebound of MAPK pathway activity, driven by enhanced wildtype KRAS signaling or receptor tyrosine kinase (RTK) signaling, in resistance to KRAS^G12C^ inhibitors^17,24^; however, such mechanisms are likely not dominant drivers in the context of MRTX1133 resistance.

**Figure 2:**
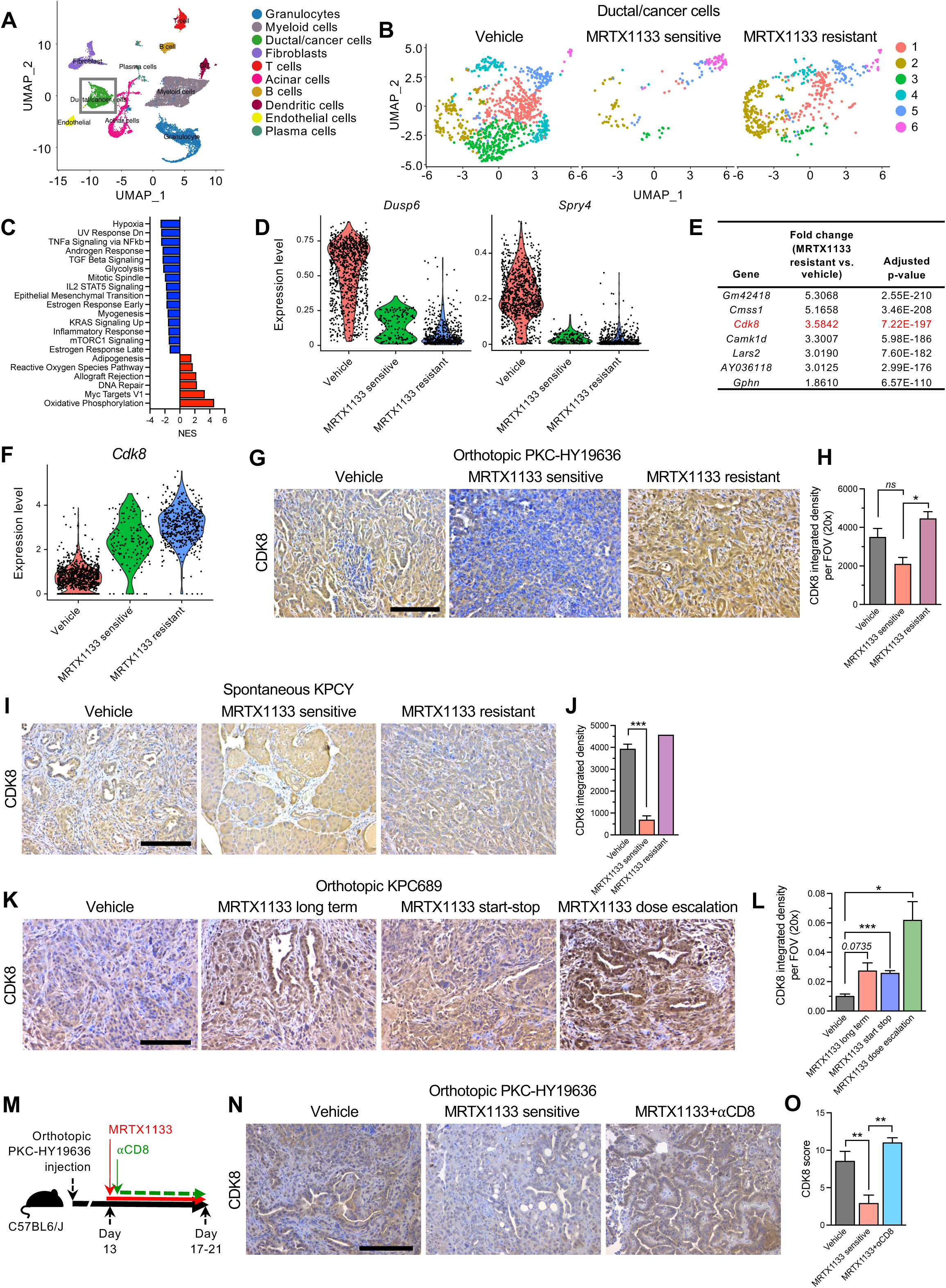
CDK8 is upregulated in MRTX1133 resistant cancer cells. (**A**) UMAP of orthotopic PKC-HY19636 tumors determined by scRNA-seq. Pancreata/tumors from 2-3 mice pooled per group. (**B**) UMAP of ductal/cancer cells of orthotopic PKC-HY19636 tumors determined by scRNA-seq. (**C**) Enriched pathways in ductal/cancer cells of MRTX1133 resistant tumors compared to vehicle orthotopic PKC-HY19636 tumors. (**D**) Violin plots of *Dusp6* and *Spry4* in ductal/cancer cells of orthotopic PKC-HY19636 tumors. (**E**) Significantly upregulated genes in ductal/cancer cells of MRTX1133 resistant tumors compared to vehicle orthotopic PKC-HY19636 tumors. (**F**) Violin plot of *Cdk8* in ductal/cancer cells of orthotopic PKC-HY19636 tumors. (**G-H**) Representative CDK8 staining (**G**) and quantification (**H**) of orthotopic PKC-HY19636 tumors. Vehicle, n=6; MRTX1133 sensitive, n=4; MRTX1133 resistant, n=3. (**I-J**) Representative CDK8 staining (**I**) and quantification (**J**) of spontaneous KPCY GEMM tumors. Vehicle, n=3; MRTX1133 sensitive, n=3; MRTX1133 resistant, n=1. (**K-L**) Representative CDK8 staining (**K**) and quantification (**L**) of orthotopic KPC689 tumors. Vehicle: n=4, MRTX1133 long term: n=5, MRTX1133 start-stop: n=4, MRTX1133 dose escalation: n=5. (**M**) Schematic representation of orthotopic PKC-HY19636 tumors treated with MRTX1133 for 5-9 days. (**N-O**) Representative CDK8 staining (**N**) and quantification (**O**) of orthotopic PKC-HY19636 tumors. Vehicle, n=7 mice; MRTX1133 sensitive, n=4 mice; MRTX1133+⍰CD8, n=4 mice. One-way ANOVA with Dunnett’s multiple comparisons test performed in (H) and (O). Two-tailed unpaired t-test performed in (J). Brown-Forsythe and Welch ANOVA with Dunnett T3’s multiple comparisons test performed in (L). Scale bars, 100 μm. Data are presented as mean ± s.e.m. except for in (D) and (F) where it is presented as violin plots. * P < 0.05, ** P < 0.01, *** P < 0.001, *ns*: not significant, or exact p-values reported.

Whole-exome sequencing (WES) was performed to further understand the genetic alterations associated with resistance to MRTX1133. MRTX1133 resistant tumors displayed heterogeneous copy number variation (CNV) patterns, with amplifications detected in *Cdk6*, *Abcb1a*, *Cdk8*, *Yap1*, and *Myc* (**Supplemental Fig. 5A**). In addition, two out of three tumors showed amplification of *Kras* (**Supplemental Fig. 5A**). In contrast to previous reports with KRAS^G12C^ inhibitors^24,25^, development of *Nras* or *Hras* mutations or amplifications was not observed in resistant tumors (**Supplemental Fig. 5A**). Further analysis of cancer cells in PKC-HY19636 orthotopic tumors by scRNA-seq validated *Cdk8* as one of the top genes upregulated in MRTX1133 resistant cancer cells (**Fig. 2E-F**). Upregulation of *Cdk8* was present in all ductal/cancer cell clusters (**Supplemental Fig. 5B**), suggesting that it is not limited to a subset of cancer cells. Immunohistochemistry for CDK8 showed that CDK8 was suppressed in MRTX1133 sensitive tumors and upregulated in MRTX1133 resistant PKC-HY19636 and KPCY tumors (**Fig. 2G-L**). Interestingly, the MRTX1133 dose escalation group showed increased expression of CDK8 in orthotopic KPC689 tumors (**Fig. 2K-L**), suggesting that higher dosages of MRTX1133 may augment the selection for treatment resistant cells with high expression of CDK8. Patient derived xenografts (PDXs) that were responsive to MRTX1133 or developed resistance to MRTX1133 or the KRAS^G12C^ inhibitor adagrasib were also evaluated for CDK8 expression. PDXs that were responsive to MRTX1133 also showed a downregulation of CDK8 with treatment, whereas resistant tumors did not (**Supplemental Fig. 5C-D**).

To further evaluate resistance to MRTX1133 treatment, an in vivo resistant PKC-HY19636 cell line was generated. The in vivo resistant cell line demonstrated diminished suppression of pERK in response to MRTX1133 compared to parental cells (approximately 2.5-fold increase in pERK IC50, **Supplemental Fig. 6A-D**). Proliferation of PKC-HY19636 in vivo resistant cells in response to MRTX1133 was also increased compared to PKC-HY19636 parental (approximately 1.6-fold increase in proliferation IC50, **Supplemental Fig. 6E-F**), further supporting resistance to MRTX1133. In addition, in vitro resistant cell lines (HY19636 and KPC689 in vitro resistant) were established by exposing cells to increasing concentrations of MRTX1133 (**Supplemental Fig. 7A**). Similar to the in vivo resistant line, the in vitro resistant cells showed reduced sensitivity to MRTX1133 and similar levels of RAS^G12D^ expression compared to parental cells (**Supplemental Fig. 7B-E**). The *Kras^G12D^* mutation was detected in parental and in vitro resistant KPC689 and PKC-HY19636 cell lines (**Supplemental Fig. 7F**). Whole-exome sequencing of in vitro resistant and parental cells revealed amplification of *Cdk6*, *Abcb1a*, *Kras*, *Yap1*, *Myc*, and *Cdk8* in resistant cells (**Supplemental Fig. 7G-H**), which may contribute to the observed increased transcriptional expression of *Cdk8*. Collectively, our data demonstrates that resistance to KRAS* inhibition is a ubiquitous phenomenon and is associated with upregulation of CDK8 across multiple PDAC models.

In order to evaluate the impact of MRTX1133 in a preventative context, we treated *LSL-Kras^G12D/+^; Pdx1-Cre* (KC) mice, which develop spontaneous PanIN precursor lesions but typically do not progress to PDAC until 12-15 months, with MRTX1133 for up to 18 weeks.^5^ MRTX1133 treatment reduced PanIN lesions, as quantified by histological evaluation and CK19 staining, and pERK abundance (**Supplemental Fig. 8A-E**). In contrast to orthotopic PKC-HY19636 tumors and spontaneous KPC GEMMs, KC mice do not present with resistance even upon long-term MRTX1133 treatment. CDK8 is not elevated in the pancreata of MRTX1133 treated mice (**Supplemental Fig. 8B, 8F**) and a reduced spatial association of CD4^+^ and CD8^+^ memory/ stem-like T cells with *Cdk8* high ductal/cancer cells was observed (**Supplemental Fig. 8G-H**), indicating a potential role for the TME in preventing the emergence of precursor lesions in KC mice.

We next determined the contribution of immune cells to the emergence of CDK8^+^ resistant cancer cells. KPC689 cells were implanted into immunocompetent C57BL/6J mice and immunodeficient NSG mice and treated with MRTX1133 for 12-13 days (**Supplemental Fig. 9A**). KPC689 tumors treated with MRTX1133 demonstrated reduced pERK in immunocompetent and immunodeficient mice (**Supplemental Fig. 9B-C**), indicating effective inhibition of KRAS* signaling in both contexts. In contrast, KPC689 tumors in NSG mice had increased CDK8 expression compared to C57BL/6J mice (**Supplemental Fig. 9B, D**). NSG mice are deficient in functional T cells, B cells, and NK cells^26^; thus, we employed antibody-mediated depletion to specifically evaluate the functional contribution of CD8^+^ T cells in restraining the expansion of resistant cells in orthotopic PKC-HY19636 tumors (**Fig. 2M**). Like NSG mice, MRTX1133 treatment of CD8^+^ T cell depleted tumors showed similar downregulation of pERK compared to CD8^+^ T cell intact tumors (**Supplemental Fig. 9E-F**), and increased CDK8 in CD8^+^ depleted tumors (**Fig. 2N-O**). We further evaluated MYC expression, which was identified as being amplified in a subset of MRTX1133 resistant tumors (**Supplemental Fig. 5A**), and observed a negligible difference in MYC in CD8^+^ depleted tumors (**Supplemental Fig. 9G-H**). Together, these data indicate that CD8^+^ T cells are critical for preventing the emergence of CDK8-expressing resistant cancer cells.

### Resistance to KRAS* inhibition is associated with reprogramming of the tumor microenvironment and lack of immunological memory response

To understand how CDK8 upregulation could alter the tumor microenvironment and potentially promote the progression of resistant tumors, spatial transcriptomics analysis and CODEX multiplex imaging of PKC-HY19636 tumors were performed. Spatial transcriptomics further revealed the distribution of stromal cells relative to ductal/cancer cells in the context of early response to MRTX1133 and MRTX1133 resistance (**Supplemental Fig. 10A-B, Supplemental Fig. 3A-C**). MRTX1133 sensitive tumors displayed reduced spatial association of ductal/cancer cells with acinar cells, endothelial cells, and Th17 cells (**Supplemental Fig. 10A-B**). In contrast, increased association of ductal/cancer cells with memory/stem-like CD8 T cells, Th2 cells, and Tregs was observed in MRTX1133 resistant tumors (**Supplemental Fig. 10A-B**), suggesting that resistance is associated with altered interactions between cancer cells and remodel the TME to promote immune evasion.

*Cdk8* high cancer cells in MRTX1133 sensitive and resistant tumors were spatially associated with immune infiltrates in the TME (**Fig. 3A-B**). Myeloid cells, dendritic cells (DCs), granulocytes, plasma cells, and Tregs displayed increased association with *Cdk8* high cancer cells compared to *Cdk8* low cancer cells (**Fig. 3A-B**), indicating that KRAS* inhibitor resistant cancer cells may remodel the local tumor microenvironment and promote immunosuppression. CODEX multiplex imaging of PKC-HY19636 tumors revealed a shift to increased relative and overall abundance of CD3^+^, CD11c^+^, and CD19^+^ cells in MRTX1133 sensitive tumors (**Fig. 3C-E**) compared to vehicle. A paucity of T cells, B cells and putative antigen-presenting cells (APCs) were observed in MRTX1133 resistant tumors potentially indicating a lack of robust immunological memory response to KRAS* inhibition (**Fig. 3C-E**). To further validate the reversal of T cell infiltrates, we examined CD8^+^ T cells in the PDAC of MRTX1133 sensitive and resistant spontaneous KPCY tumors (**Supplemental Fig. 10C-D**). Consistent with our findings from the orthotopic PKC-HY19636 tumors, the KPCY tumors demonstrated paucity of CD8^+^ T cell infiltrates in MRTX1133 resistant tumors (**Supplemental Fig. 10C-D**). We validated alterations in the TME using an alternative resistance model, wherein in vitro parental and MRTX1133 resistant KPC689 cell lines were implanted and treated with MRTX1133 (**Supplemental Fig. 11A**). In vitro resistant KPC689 cells were established by exposing the parental line to high concentration of MRTX1133 (**Supplemental Fig. 7A**). The resistant and parental orthotopic tumors were treated with MRTX1133 following establishment of baseline PDAC (**Supplemental Fig. 11A-B**). Analysis of KPC689 resistant and parental tumors demonstrate an increased frequency of myeloid cells and exhausted CD8^+^ T cells, accompanied by a decrease in B cell, CD8^+^ effector T cell, CD4^+^ T cell, and CD8^+^ memory/stem-like T cell frequencies in the MRTX1133 resistant tumors (**Supplemental Fig. 11G-J**), suggesting a lack of a CD8^+^ T cell memory response. Taken together, our data suggests that resistance to KRAS* inhibition is associated with an upregulation of CDK8 in cancer cells and a reversal of T and B cell infiltration from the MRTX1133 sensitive tumors suggesting that CDK8 and its downstream effectors could be potential mediators of MRTX1133 resistance.

**Figure 3:**
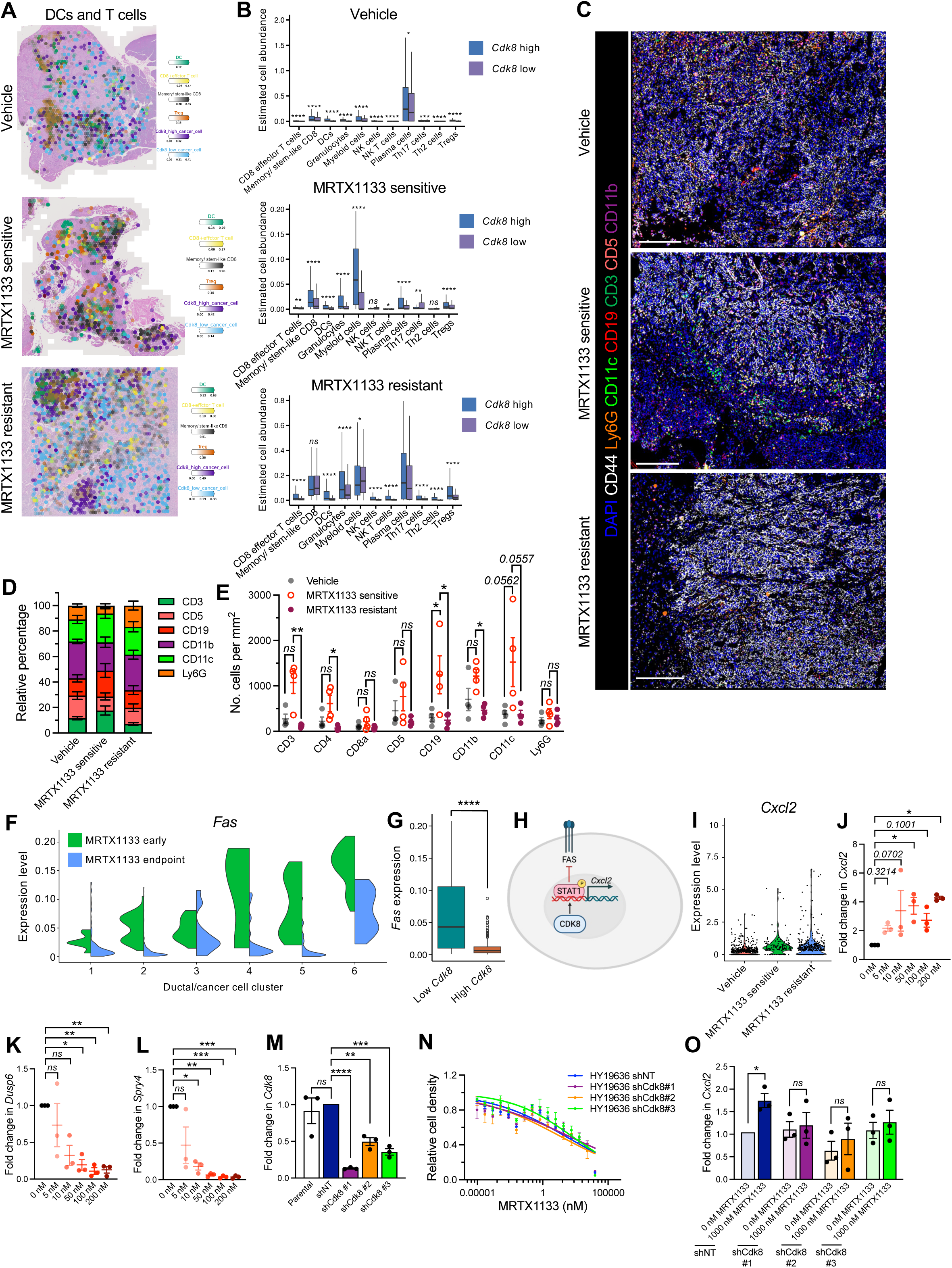
MRTX1133 resistance alters the tumor microenvironment and leads to downregulation of FAS and upregulation of CDK8 and CXCL2. (**A-B**) Spatial transcriptomics representative images (**A**) and quantification (**B**) of orthotopic PKC-HY19636 tumors. Spots were stratified by *Cdk8* high and low areas. Vehicle, n=3; MRTX1133 sensitive, n=3; MRTX1133 resistant, n=3. (**C-E**) Representative CODEX images (**C**), quantification of relative percentages of immune cell types (**D**), and number of cells per tissue area (**E**) of PKC-HY19636 orthotopic tumors. Vehicle, n=4; MRTX1133 sensitive, n=4; MRTX1133 resistant, n=4. (**F**) *Fas* expression stratified by *Cdk8* expression level in ductal/cancer cells of MRTX1133 sensitive (early) and resistant (endpoint) orthotopic PKC-HY19636 tumors evaluated by scRNA-seq. (**G**) *Fas* expression stratified by *Cdk8* expression level in ductal/cancer cells of MRTX1133 sensitive and resistant orthotopic PKC-HY19636 tumors evaluated by scRNA-seq. (**H**) Schematic of CDK8 signaling. Created in BioRender. McAndrews, K. (2025) https://BioRender.com/1xttcgt. (**I**) Violin plot of *Cxcl2* expression in ductal/cancer cells of PKC-HY19636 tumors. (**J**) *Cxcl2* mRNA expression in KPC689 cells treated with varying concentrations of MRTX1133. n=3 independent experiments. (**K-L**) *Dusp6* (**K**) and *Spry4* (**L**) mRNA expression in KPC689 cells treated with varying concentrations of MRTX1133 for 6 hours. Data is presented relative to *Gapdh* C_T_ values and 0 nM MRTX1133. n=3 independent experiments. (**M**) mRNA expression of *Cdk8* in parental, shNon-targeting (shNT), and shCdk8 HY19636 cells. Data are normalized to *18s* C_T_ values and shNT. n=3 independent experiments. (**N**) Relative cell density of HY19636 cells treated with the indicated concentrations of MRTX1133 for 3 days. (**O**) mRNA expression of *Cxcl2* following MRTX1133 treatment for 24 hours. Data are normalized to *18s* C_T_ values and shNT. n=3 independent experiments. Data are presented as mean ± s.e.m. except for in (B), (F), (G), and (I) where it is presented as median and range in (B, G) and violin plots in (F, I). Two-sided Mann-Whitney test was performed in (B) and (G). Kruskal-Wallis test with Dunn’s multiple comparisons test performed for CD3, CD4, and CD5 in (E) and one-way ANOVA with Dunnett’s multiple comparisons test performed for CD8, CD19, CD11b, CD11c, and Ly6G in (E). Scale bars, 250 μm. One-way ANOVA with Dunnett’s multiple comparison test performed in (J), (K), (L), and (M). Unpaired two-tailed t-test performed in (O). * P < 0.05, ** P < 0.01, *** P < 0.001, **** P < 0.0001, *ns*: not significant.

### KRAS* inhibitor resistance leads to downregulation of FAS and upregulation of CDK8 and CXCL2

Next, we probed whether CDK8 upregulation abrogates KRAS* inhibitor-mediated cell death in the PDAC. Our previous work showed that FAS expression on cancer cells induced by KRAS* inhibition promotes the induction of apoptosis through its engagement with FASL on CD8^+^ T cells.^3,10^ Analysis of *Fas* expression in ductal/cancer cells of PKC-HY19636 tumors by scRNA-seq revealed reduced expression in MRTX1133 resistant cancer cells (**Fig. 3F**). In addition, stratifying ductal/cancer cells based on *Cdk8* expression revealed an inverse association between *Cdk8* expression and *Fas* expression (**Fig. 3G, Supplemental Fig. 12A**), suggesting a potential mechanism by which resistant cancer cells evade immune recognition and clearance.

Previous studies have demonstrated that CDK8 acts to phosphorylate STAT1, which can inhibit expression of the death receptor FAS^27^ and induce CXCL2 expression^28^ (**Fig. 3H**). Increased *Cxcl2*, a downstream target of CDK8^28^, was observed in ductal/cancer cells of MRTX1133 resistant tumors (**Fig. 3I**). Granulocytes and myeloid cells exhibit high *Cxcl2* expression which is not altered in MRTX1133 resistant tumors (**Supplemental Fig. 12B**). Increased *Cxcl2* expression is present in acinar cells and DCs of resistant tumors (**Supplemental Fig. 12B**), which may in conjunction with cancer cell-derived CXCL2 contribute to altered immune infiltration. Analysis of other ligands that bind to CXCR2, the cognate receptor for CXCL2, revealed minimal expression of *Cxcl1*, *Cxcl3*, and *Cxcl5* (**Supplemental Fig. 12C**), suggesting that CXCL2 is the primary ligand produced by MRTX1133 resistant cancer cells. In vitro treatment of KPC689 cells with MRTX1133 led to upregulation of *Cxcl2* in a dose-dependent manner (**Fig. 3J**). In contrast, the KRAS* downstream transcriptional targets *Dusp6* and *Spry4* were inhibited at 6 hours post-treatment with MRTX1133 (**Fig. 3K-L**), demonstrating continued suppression of KRAS* signaling. Next, to determine the functional impact of CDK8 on *Cxcl2* expression, we generated PKC-HY19636 cells with stable knockdown of *Cdk8* (**Fig. 3M**). Genetic suppression of *Cdk8* in PKC-HY19636 cells did not impact the proliferative response to MRTX1133 (**Fig. 3N**); however, the MRTX1133 treatment associated increase in *Cxcl2* expression was abrogated in shCdk8 cells (**Fig. 3O**). Together, these data suggest that CDK8 does not play a significant role in cancer cell proliferation in response to MRTX1133, rather impacts cancer cell extrinsic signaling, and indicate that *Cxcl2* is a downstream target of CDK8.

CXCL2 binds to CXCR2 expressed on neutrophils and myeloid cells and inhibition of CXCR2 suppresses PDAC metastasis and increases T cell infiltration to sensitize PDAC to anti-PD-1 ICB^29^. Myeloid subsets of PKC-HY19636 tumors were further analyzed by scRNA-seq, where 5 myeloid clusters were defined. Cells in cluster 1 express genes associated with myeloid derived suppressor cells (MDSCs; *Arg1*, *Vegfa*, *Thbs1*, and *Spp1*), cluster 2 express genes associated with ApoE and lipid metabolism (*Apoe*, *C1q*, *Cd63*, and *Cd81*), cluster 3 express antigen presentation genes (*H2-Eb1*, *H2-Aa*, *H2-Ab1*, and *Cd74*), cluster 4 express genes associated with alternative activation (*Mrc1*, *Cd163*, and *Folr2*), and cluster 5 express proliferation genes (*Top2a*, *Mki67*, and *Stmn1*) (**Supplemental Fig. 12D**). MRTX1133 sensitive tumors demonstrate a decrease in relative abundance of cluster 1 and an increase in the relative abundance of cluster 2, cluster 3, and cluster 4 compared to vehicle and MRTX1133 resistant tumors (**Supplemental Fig. 12E**). Previous studies have implicated MDSCs in generating an immunosuppressive microenvironment characterized by reduced T cell infiltration and increased T cell exhaustion^29,30^. In addition, regulatory T cells can shape the expansion and differentiation of MDSCs in the context of inflammation^31^, indicating that bidirectional signaling between T cells and MDSCs may occur. This suggests a shift from predominantly myeloid cells that promote antigen presentation and anti-tumor immunity to immunosuppressive myeloid cells as resistance to MRTX1133 emerges.

### CDK8 and CXCR2 inhibition overcome KRAS* inhibitor resistance

The observed upregulation of CDK8 and CXCL2 in KRAS* inhibitor resistant cancer cells suggested that they may be targets to prevent resistance. Orthotopic PKC-HY19636 tumors were treated with a CDK8 inhibitor (CDK8i, BI1347) or an inhibitor of the receptor for CXCL2, CXCR2 (CXCR2i, SB225002), alone or in combination with MRTX1133 and had no impact on bodyweight during treatment (**Fig. 4A, Supplemental Fig. 13A**). CDK8i or CXCR2i as single agents had no impact on overall survival; however, when combined with MRTX1133, a significant increase in survival was observed with both the inhibitors (**Fig. 4B**). At one-week post-treatment, the weights of tumor/pancreas treated with MRTX1133 were similar to MRTX1133 combined with CDK8i or CXCR2i (**Fig. 4C**). Histological analysis further confirmed similar tumor burden with MRTX1133 single agent treatment compared to combination with CDK8i or CXCR2i, one-week post-treatment (**Fig. 4D-E**). MRTX1133 alone or in combination with CDK8i or CXCR2i was associated with reduced pERK abundance (**Fig. 4D, 4F**). In addition, decreased CDK8 was confirmed in tumors treated with CDK8i (**Fig. 4D, 4G**). CD206, a marker of myeloid derived suppressor cells (MDSCs) that express CXCR2, were significantly decreased in CXCR2i treated tumors (**Fig. 4D, 4H**). Moreover, MRTX1133 resistant tumors had increased CD206 abundance, which was abrogated with either CDK8i or CXCR2i (**Supplemental Fig. 13B-C**). These results suggest that MRTX1133 in combination with CDK8i and CXCR2i resulted in similar tumor growth inhibition response initially and the significant survival increase associated with the combination therapy (**Fig. 4B**) could likely be due to prevention of resistance to MRTX1133.

**Figure 4:**
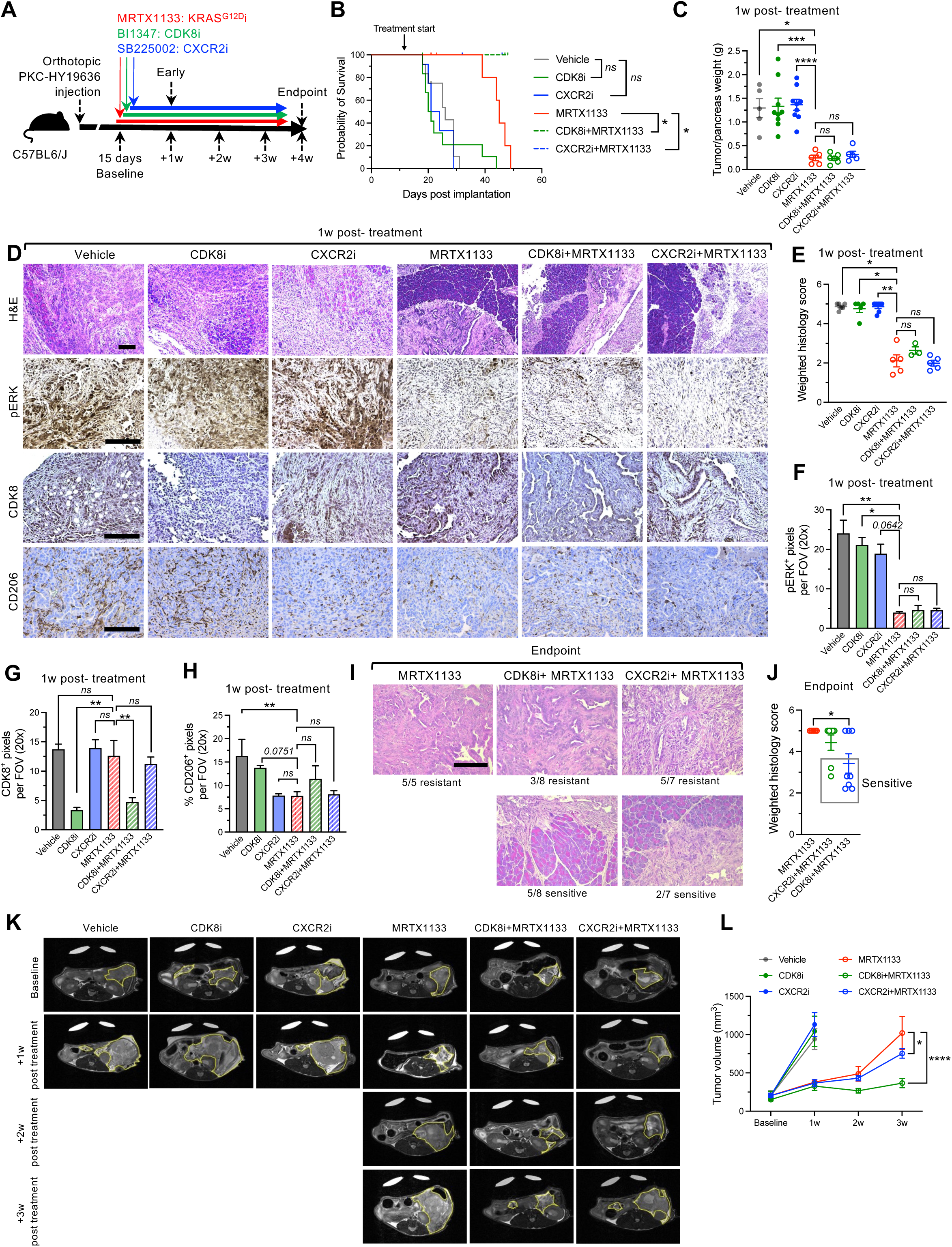
CDK8 and CXCR2 inhibition reverses MRTX1133 resistance. (**A**) Schematic of orthotopic PKC-HY19636 tumors treated with MRTX1133, CDK8i (BI1347), and CXCR2i (SB22502). (**B**) Survival curve of orthotopic PKC-HY19636 mice. Vehicle, n=12; CDK8i, n=12; CXCR2i, n=12; MRTX1133, n=12; MRTX1133+CDK8i, n=12; MRTX1133+ CXCR2i, n=12. Survival experiment was terminated shortly after all the MRTX1133 treated mice reached endpoints. (**C**) Tumor/pancreas weights at 1-week post-treatment. Vehicle, n=5; CDK8i, n=9; CXCR2i, n=9; MRTX1133, n=5; MRTX1133+CDK8i, n=6; MRTX1133+ CXCR2i, n=5. (**D**) Representative H&E, pERK, CDK8, and CD206 staining at 1-week post-treatment. (**E**) Weighted histology score. Vehicle, n=6; CDK8i, n=5; CXCR2i, n=7; MRTX1133, n=5; CDK8i+MRTX1133, n=3; CXCR2i+MRTX1133, n=5. (**F-H**) Quantification of pERK (**F**), CDK8 (**G**), and CD206 (**H**) staining. pERK: Vehicle, n=5; CDK8i, n=5; CXCR2i, n=5; MRTX1133, n=5; CDK8i+MRTX1133, n=4; CXCR2i+MRTX1133, n=5. CDK8: Vehicle, n=5; CDK8i, n=3; CXCR2i, n=4; MRTX1133, n=4; CDK8i+MRTX1133, n=4; CXCR2i+MRTX1133, n=3. CD206: Vehicle, n=5; CDK8i, n=5; CXCR2i, n=6; MRTX1133, n=5; CDK8i+MRTX1133, n=3; CXCR2i+MRTX1133, n=5. (**I-J**) Representative H&E (**I**) and weighted histology score (**E**) of murine pancreata at endpoint. MRTX1133, n=5; CDK8i+MRTX1133, n=8; CXCR2i+MRTX1133, n=7. (**K-L**) Representative images (**K**) and quantified tumor volume measured by MRI (**L**). Yellow lines demarcate tumor regions. Vehicle, n=7; CDK8i, n=7; CXCR2i, n=7; MRTX1133, n=6; CDK8i+MRTX1133, n=8; CXCR2i+MRTX1133, n=8. Gehan-Breslow-Wilcoxon test performed in (B). Brown-Forsythe and Welch ANOVA with Dunnett’s T3 multiple comparisons test performed in (C). One-way ANOVA with Dunnett’s multiple comparisons test performed in (G) and (H). Kruskal-Wallis with Dunn’s multiple comparisons test performed in (E) and (F). Two-way ANOVA with Dunnett’s multiple comparisons test performed in (L). Scale bars, 100 μm. Data are presented as mean ± s.e.m. * P < 0.05, ** P < 0.01, *** P < 0.001, **** P < 0.0001, *ns*: not significant.

To assess the impact of CDK8i and CXCR2i in preventing resistance to MRTX1133, we examined the histology of the tumors at endpoint (**Fig. 4I-J**). At endpoint, all the MRTX1133 treated mice (100%) developed resistance, whereas 5/8 (∼63%) of MRTX1133+CDK8i and 2/7 (28%) of the MRTX1133+ CXCR2i treated mice demonstrated tumor growth inhibition. MRI analysis revealed that MRTX1133 initially controlled tumor growth, but long-term treatment led to tumor relapse (**Fig. 4K-L, Supplemental Fig. 13D**). In contrast, MRTX1133 combined with CDK8i or CXCR2i led to more durable responses, with reduced tumor burden (**Fig. 4K-L, Supplemental Fig. 13D**). Our data demonstrates that upregulation of CDK8 functionally contributes to resistance to MRTX1133 and pharmacological targeting of CDK8 and its downstream target CXCL2 impedes the development of resistance to KRAS* inhibition in PDAC.

### Priming of KRAS* inhibitor resistant tumors with CDK8 inhibition sensitizes tumors to αCTLA-4 checkpoint blockade immunotherapy

We next probed the contribution of the adaptive immune response in reversing resistance to KRAS* inhibition in combination with CDK8 inhibition and checkpoint immunotherapy. Analysis of antigen presentation related genes in the ductal/cancer cells of PKC-HY19636 tumors revealed increased expression in MRTX1133 sensitive tumors compared to vehicle; however, expression was downregulated or unchanged in MRTX1133 resistant tumors (**Fig. 5A**). CODEX multiplex imaging of PKC-HY19636 tumors revealed an increase in CD3^+^TCRβ^+^ and CD4^+^TCRβ^+^ cells in MRTX1133 sensitive tumors, which was reduced in MRTX1133 resistant tumors (**Fig. 5B-C**). Similarly, increased abundance of putative antigen presenting cells (APCs, CD11b^+^MHCII^+^ and CD11c^+^MHCII^+^) was observed in MRTX1133 sensitive tumors compared to MRTX1133 resistant tumors (**Supplemental Fig. 13E-F**), indicating a loss of antigen presentation in the context of resistance. This finding suggested that resistant tumors would not be responsive to checkpoint blockade, and that CDK8 inhibition could potentially remodel the immune microenvironment and sensitize MRTX1133 resistant tumors to ICB. Our prior work with the orthotopic PKC-HY19636 model demonstrated that upfront combination of αCTLA-4 with the MRTX1133 therapy following establishment of advanced tumors resulted in robust tumor inhibition response^10^. In a second model, MRTX1133 in upfront combination with αCTLA-4 demonstrated synergy and long-term survival in KPC-T mice (**Fig. 5D-E, Supplementary Fig. 13G-H**). Our results establish that early CDK8 inhibition or αCTLA-4 ICB synergizes with KRAS* inhibition to impede development of resistance to MRTX1133 therapy. Further analysis of FAS expression on cancer cells and T cell proliferation during the development of MRTX1133 resistance revealed that Fas expressing cancer cells decreased in MRTX1133 resistant tumors (**Fig. 5F-H**). Although addition of αCTLA-4 resulted in an increase in FAS^+^ CDK8^−^ cancer cells, a lack of CD8^+^ T cell infiltration in the MRTX1133 resistant tumors likely resulted in persistence of the FAS expressing cancer cells which require CD8^+^ T cells for their elimination (**Fig. 5F-H, Supplemental Fig. 14A-C**). Collectively, our results indicate that the addition of αCTLA-4 or CDK8i simultaneously with MRTX1133 prevents the development of resistance to KRAS* inhibition.

**Figure 5:**
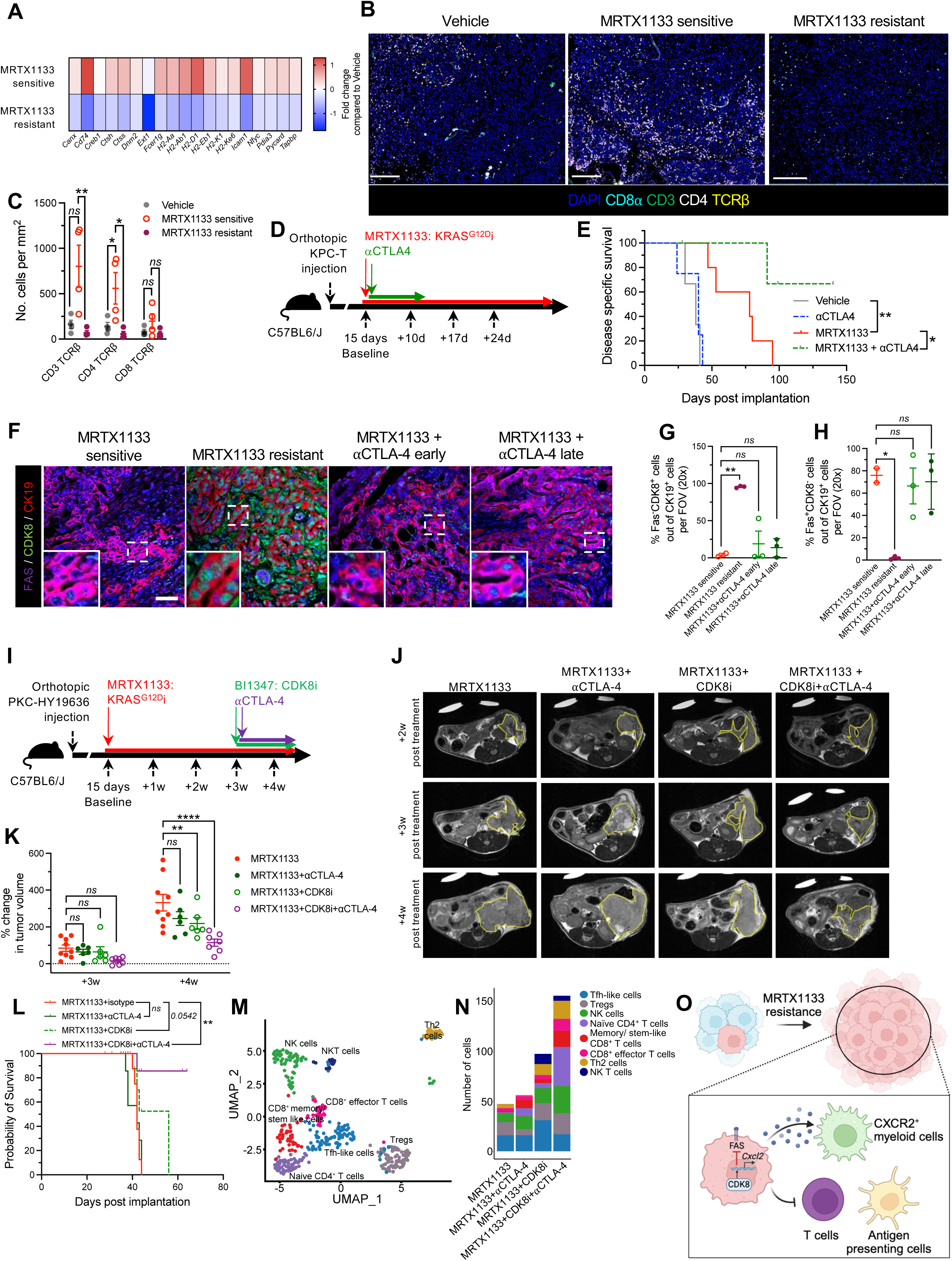
Resistance to MRTX1133 leads to loss of antigen presentation which can be reversed with inhibition of CDK8. (**A**) Expression of antigen presentation genes of ductal/cancer cells of PKC-HY19636 tumors compared to vehicle treated mice as determined by scRNA-seq analysis. (**B-C**) Representative CODEX images of CD3, CD8α, CD4, and TCRβ (**B**) and quantification of cell abundance per tissue area of PKC-HY19636 orthotopic tumors (**C**). Scale bars, 250 μm. (**D**) Schematic of orthotopic PKC-HY19636 tumors treated with MRTX1133 and αCTLA4. (**E**) Survival curve of orthotopic PKC-HY19636 mice. Vehicle, n=3; αCTLA4, n=4; MRTX1133, n=5; MRTX1133+ αCTLA4, n=6. (**F-H**) Representative images (**F**) and quantification of FAS^−^CDK8^+^CK19^+^ (**G**) and FAS^+^CDK8^−^CK19^+^ (**H**) cells in orthotopic PKC-HY19636 tumors. MRTX1133 sensitive, n=2; MRTX1133 resistant, n=3; MRTX1133+⍰CTLA-4 early, n=3; MRTX1133+⍰CTLA-4 late, n=3. Scale bar, 50 μm. (**I**) Schematic of orthotopic PKC-HY19636 tumors treated with MRTX1133, CDK8i (BI1347), and αCTLA-4. (**J-K**) Representative images (**J**) and quantified tumor volume measured by MRI relative to tumor volume at 2 weeks following MRTX1133 treatment initiation (**K**). Yellow lines demarcate tumor regions. MRTX1133, n=9; MRTX1133+αCTLA-4, n=6; MRTX1133+CDK8i, n=7; MRTX1133+CDK8i+αCTLA-4, n=7. (**L**) Survival curve of orthotopic PKC-HY19636 mice. MRTX1133, n=10; MRTX1133+αCTLA-4, n=8; MRTX1133+CDK8i, n=11; MRTX1133+CDK8i+αCTLA-4, n=10. (**M**) UMAP of T and NK cells of orthotopic PKC-HY19636 tumors determined by scRNA-seq. Pancreata/tumors from 3 mice pooled per group. (**N**) Overall abundance of T and NK cell subsets in PKC-HY19636 determined by scRNA-seq. (**O**) Graphical abstract. Created in BioRender. McAndrews, K. (2025) https://BioRender.com/awmhuoy. Kruskal-Wallis with Dunn’s multiple comparisons test performed for CD3 TCRβ in (C), one-way ANOVA with Dunnett’s multiple comparisons test performed for CD4 TCRβ and CD8 TCRβ in (C), and log-rank test performed in (E and L). One-way ANOVA with Dunnett’s multiple comparisons test performed in (G), (H). Dunnett’s multiple comparisons test performed in (K). * P < 0.05, ** P < 0.01, **** P < 0.001, *ns*: not significant.

Next, to evaluate whether priming tumors with CDK8 inhibition could reverse resistance to KRAS* inhibition through eliciting T cell infiltration and sensitize tumors to ICB, PKC-HY19636 orthotopic tumor bearing mice were treated with MRTX1133 for 3 weeks to establish resistance (**Fig. 5I**). Treatment with ⍰CTLA-4 and CDK8i was initiated at 3 weeks (**Fig. 5I**). Combination of MRTX1133 with ⍰CTLA-4 at the time of development of resistance did not impede tumor growth likely due to the absence of T cell infiltrates in MRTX1133 resistant tumors (**Fig. 5B-C, Supplemental Fig. 14A-B**, **Fig. 5I-L**). Although CDK8i impeded development of resistance to MRT1133 (vide supra-Fig. **4**), addition of CDK8i in MRTX1133 resistant tumors resulted in a marginal improvement in survival (**Fig. 5J-L**). CDK8i in combination with ⍰CTLA-4 resulted in robust inhibition of tumor growth with prolonged survival (**Fig. 5J-L**). Combination of CDK8i and ⍰CTLA-4 reversed MRTX1133 resistance likely indicating that CDK8i primed the tumor microenvironment to ICB (**Fig. 5M-N**). Analysis of expression of immune checkpoint markers indicated that *Ctla4* is the primary immune checkpoint expressed in PKC-HY19636 tumors (**Supplemental Fig. 14D**). In contrast to the combination with ⍰CTLA-4, CDK8i with PD-1 did not significantly impact the emergence of MRTX1133 resistance (**Supplemental Fig. 14E-G**), potentially due to the presence of effector regulatory T cells that suppress CD8^+^ T cells^32^.

Further, scRNA seq analysis of the tumor infiltrating T and NK cell populations demonstrated an increase in the memory/stem-like CD8^+^ T cells, CD8^+^ effector T cells, naïve CD4^+^ T cells, and NK cells in the tumors of CDK8i and ⍰CTLA4 combination treated mice (**Fig. 5M-N, Supplementary Fig. 15A**). The myeloid compartment of tumors treated with MRTX1133 and CDK8i was analyzed by scRNA-seq and a decrease in the relative abundance of myeloid cells was observed (**Supplemental Fig. 15B-C**). Further subclassification of myeloid cells revealed minor changes in the relative abundance of myeloid subpopulations (**Supplemental Fig. 14D-E**), suggesting that CDK8 inhibition likely broadly targets myeloid cells.

Collectively, our data establishes that long-term treatment with a pharmacological inhibitor of KRAS*, MRTX1133, resulted in resistance leading to mice succumbing to PDAC. Upregulation of CDK8 in the pancreatic cancer cells contributes to development of resistance to MRTX1133 therapy and inhibition of CDK8 and the downstream target CXCR2 impedes development of resistance. MRTX1133 treatment acutely increased the T cell infiltration in PDAC, rendering the tumors sensitive to αCTLA-4 ICB. However, lack of long-term immunological memory following MRTX1133 therapy resulted in paucity of T cells and APCs in resistant tumors (**Fig. 5O**). Although ICB did not reverse resistance to MRTX1133, combination of CDK8 inhibition rendered MRTX1133 resistant PDAC amenable to αCTLA-4 ICB.

## Discussion

Small molecule inhibitors of KRAS* have demonstrated efficacy in controlling PDAC in both preclinical models and in patients;^9–11,19,33–36^ however, acquired resistance has been reported in the context of KRAS^G12C^ inhibition.^18,25,37^ Previous studies have identified feedback activation of EGFR as promoting MRTX1133 resistance in pancreatic and colorectal cancer^38,39^ similar to what has been reported with KRAS^G12C^ inhibitors^17^; however, it is unclear whether resistance mechanisms unique to KRAS^G12D^ inhibition exist given the distinct impacts KRAS mutations have on pancreatic cancer initiation and progression.^20^ Inherent differences in the non-covalent binding of KRAS^G12D^ inhibitors compared to covalent binding KRAS^G12C^ inhibitors may contribute to mechanisms of inhibition and emergence of resistance. KRAS^G12C^ inhibitors bind the GDP-bound inactive state^21,40,41^ whereas the KRAS^G12D^ inhibitor MRTX1133 is capable of binding both the inactive and active states,^11^ raising the possibility that specificity of inhibitor binding may impact the mechanisms of resistance that emerge. Moreover, KC mice (with no defects in tumor suppressor genes) did not develop resistance to MRTX1133, in comparison to PDAC models with loss of or gain of function mutant p53. This suggests that tumor suppressors may contribute to resistance to KRAS* inhibitors. Indeed, CRISPR screening identified *TP53*, *RB1*, and *KEAP1* as mediators of partial resistance to MRTX1133.^11^ Mutations in *KRAS*, *NRAS*, *MRAS*, *RET*, *MAP2K1*, *BRAF*, *NF1*, and *PTEN*, amplifications in *KRAS^G12C^* and *MET*, and oncogenic fusions have been described in tumors resistant to KRAS^G12C^ inhibitors^18,25^ and *Myc* amplification has been identified in the context of resistance to pan RAS inhibitors^42^. Moreover, feedback signaling through wildtype KRAS and RTKs can drive adaptive resistance to KRAS^G12C^ inhibition, although lineage specificity of such signaling has been reported^17,24^. Amplifications in *Cdk6*, *Abcb1a*, *Kras*, *Yap1*, and *Myc* were identified in MRTX1133 resistant tumors in our study, and more in-depth analysis of the genomic landscape of resistant murine and patient tumors will provide critical insight into the contribution of genetic alterations to RAS inhibitor resistance and identify additional therapeutic vulnerabilities. Earlier studies have implicated YAP/TAZ, CDK4/6, and MYC as potential mediators of resistance to KRAS^G12D^ or MEK inhibition ^14,43,44^.

Here, we identify CDK8 as promoting resistance to KRAS^G12D^ inhibitor, with a focus on its impact on the TME. CDK8 promotes resistance in part through suppression of FAS expression and increased CXCL2 expression, which generates an immunosuppressive microenvironment. In the context of cancer, CDK8 has been shown to be an oncoprotein that promotes transcription,^45,46^ suggesting that CDK8 may alter the expression of additional genes outside of *Fas* and *Cxcl2* to impact cancer cell intrinsic and extrinsic processes. Evaluation of such potential transcriptional changes induced by CDK8 may provide additional insights into the mechanisms by which it promotes resistance and will be important for future studies. It is likely that co-targeting of CXCL2/CXCR2 with CDK8i would be redundant as CXCL2 is downstream of CDK8 and redundancy in cytokine receptors^47^ may explain the reduced therapeutic effectiveness of targeting CXCR2 compared to CDK8 that was observed. Based on our data, the combination of CDK8i with ICB is expected to provide more durable responses. In addition, the development of more specific CDK8 small molecule inhibitors with improved toxicity and pharmacokinetic profiles would enable clinical translation. While clinical development of MRTX1133 has been terminated due to formulation challenges (ClinicalTrials.gov, NCT05737706), future studies evaluating alternate KRAS* targeting strategies will clarify whether CDK8 upregulation is a conserved mechanism of resistance and common therapeutic vulnerability.

*Camk1d* was also identified as upregulated in MRTX1133 resistant cancer cells and has previously been associated with expression of FAS in multiple myeloma cancer cells and is coregulated with PD-L1 expression to potentially mediate resistance to anti-PD-1 therapy^48^, suggesting that it may be another potential mediator of immune modulation in the context of resistance and may be important target for future studies. Previous studies employing inducible models of *Kras^G12D^* genetic extinction identified HDAC5 as promoting CCL2 expression to increase myeloid infiltration and promote the relapse of tumors with *Kras^G12D^* extinguished compared to *Kras^G12D^* intact tumors^16^. We did not specifically evaluate the contribution of HDAC5 and CCL2 in this study and if the mechanisms identified through genetic suppression of KRAS are conserved with pharmacological KRAS inhibition are still being unraveled and will be important for future studies.

We demonstrate that MRTX1133 treatment initially increases the infiltration of TCRβ^+^ T cells and CD11c^+^ cells, which are lost with prolonged MRTX1133 treatment and the emergence of resistance. This suggests that there is a narrow window of opportunity to further stimulate CD8^+^ T cell mediated killing of cancer cells with ICB to effectively clear cancer cells and prevent the expansion of resistant cancer cells. Indeed, combination of MRTX1133 with αCTLA-4 impedes the development of resistance and improves survival outcomes.^10^ After CDK8-mediated resistance emerges, cancer cells likely evade immune recognition through reduced expression of FAS, which can directly engage with FASL on CD8^+^ T cells to induce cancer cell apoptosis, increased expression of CXCL2, which can signal to myeloid cells to indirectly promote suppression of T cells, and loss of antigen presentation. CXCR2 is expressed by myeloid-derived suppressor cells which can suppress T cell responses^49,50^; thus, CXCR2 inhibition likely promotes CD8^+^ T cell activity through both FASL-dependent and independent mechanisms.

Emergence of additional mutations has been reported in tumors resistant to KRAS^G12C^ inhibition;^18,25^ however, the mutational landscape of MRTX1133 resistant tumors has not been reported and whether such mutations functionally contribute to cancer progression is unknown. The emergence of new mutations in resistant cancer cells in combination with their suppression of CD8^+^ T cells may enable cancer cell escape from immune recognition. Indeed, ICB did not show therapeutic benefit in MRTX1133 resistant tumors, and additional priming with CDK8i was needed to sensitize MRTX1133 resistant tumors to ICB.

The development of small molecule inhibitors of KRAS^G12D^ has enabled targeting the dominant driver oncogene in PDAC. While KRAS^G12D^ inhibitors demonstrate control of PDAC growth, we report with long term treatment that resistance occurs. We identify CDK8 as promoting resistance through inhibition of FAS expression and induction of CXCL2 secretion to reprogram the TME into an immunosuppressive state. Our studies provide a rationale for targeting CDK8 in resistant tumors and employing ICB prior to the emergence of resistance to KRAS^G12D^ small molecule inhibitors.

## Methods

### Cell culture

HY19636 were isolated from *Ptf1a-Cre; LSL-Kras^G12D/+^; Trp53^F/+^* (PKC) tumors^51^ and KPC689^52^ and KPC-T^22^ were isolated from *Pdx1-Cre; LSL-Kras^G12D/+^; LSL-Trp53^R172H/+^* (KPC) tumors as previously described. HY19636 in vivo resistant cells were isolated from an orthotopic HY19636 tumor that developed resistance to MRTX1133. The tumor was digested in 4 mg/mL collagenase IV (Gibco) and 4 mg/mL dispase (Gibco) in RPMI with 1% PSA for 50 minutes at 37°C with shaking, followed by consecutive filtering through 70 μm and 40 μm filters. Cells were washed with growth media: RPMI (Corning) with 10% FBS (Avantor) and 1% penicillin-streptomycin with antimycotics (PSA, Corning), centrifuged at 500g for 5 minutes, then seeded on collagen I coated plates (Sigma Aldrich) in growth media. KPC689 and HY19636 in vitro resistant cells were generated by exposing KPC689 parental and HY19636 parental cells to increasing concentrations of MRTX1133 (WuXi AppTech) over the course of 2 to 3 months: 150 nM MRTX1133, followed by 300 nM MRTX1133, 600 nM MRTX1133, and 900 nM MRTX1133. In vitro resistant cells were then maintained in 900 nM MRTX1133 unless noted otherwise. KPC689 were transduced with a GFP-luciferase lentivirus (Capital Biosciences) followed by selection with and maintenance in 2 μg/mL puromycin (Sigma Aldrich). HY19636 cells were transduced with non-targeting shRNA lentiviral particles (shNT, pLKO.1-puro Non-Mammalian shRNA Control Transduction Particles, Sigma Aldrich SHC002VN) or shCdk8 lentiviral particles (Sigma Aldrich SHCLNV, **Table 1**) with 8 μg/mL polybrene (Millipore) followed by selection with and maintenance in 8 μg/mL puromycin (Sigma Aldrich).

**Table 1:**
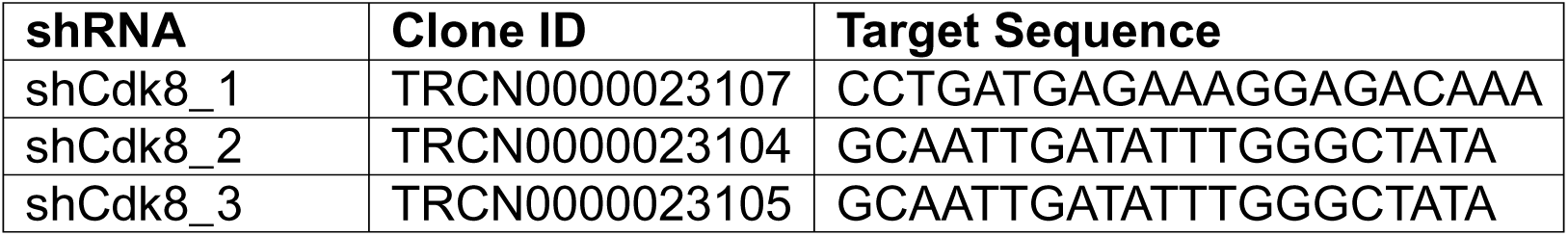
shCdk8 clones and target sequences.

Cells were cultured at 37°C and 5% CO_2_ in the media listed in **Table 2**. All cell lines were routinely tested for mycoplasma and confirmed negative with LookOut Mycoplasma PCR Detection Kit (Sigma Aldrich).

**Table 2:** Cell line origins and culture conditions.

| Cell line | Origin | Culture media |
| --- | --- | --- |
| HY19636 | <sup>51</sup> | RPMI (Corning), 10% FBS, 1% penicillin-streptomycin (PS, Corning) |
| KPC689 | <sup>52</sup> | RPMI, 10% FBS, 1% PS |
| KPC-T | <sup>22</sup> | RPMI, 10% FBS, 1% PS |
| HY19636 in vivo resistant | This manuscript | RPMI, 10% FBS, 1% PSA |
| KPC689 in vitro resistant | This manuscript | RPMI, 10% FBS, 1% PS, 900 nM MRTX1133 |
| HY19636 in vitro resistant | This manuscript | RPMI, 10% FBS, 1% PS, 900 nM MRTX1133 |

#### Animal studies

Male and female 8- to 10-week-old C57BL6/J mice were purchased from Jackson Laboratories. 5 × 10^5^ PKC-HY19636, KPC689 GFP-luciferase, KPC689 parental, KPC689 in vitro resistant, or KPC-T cells were orthotopically injected into the pancreas tail with a 27-gauge Hamilton syringe. PDX experiments were previously described.^10^ Spontaneous *LSL-Kras^G12D/+^; LSL-Trp53^R172H/+^; Pdx1-Cre; LSL-EYFP* (KPCY) mice^5,53,54^ were generated, maintained on mixed background, and enrolled in treatment at approximately 85 days when the presence of tumors was confirmed by MRI. *LSL-Kras^G12D/+^; Pdx1-Cre* (KC) were generated, maintained on mixed background, and enrolled in treatment at approximately 49 days. For orthotopic experiments, treatment was initiated at the timepoints indicated in the figures. Imaging of KPC689 GFP-luciferase tumors was performed with IVIS (Xenogen Spectrum) following injection of D-luciferin (Goldbio) according to the manufacturer’s instructions. The tumor size of PKC-HY19636, KPC689 parental, KPC689 in vitro resistant, and KPC-T orthotopic tumors and KPC GEMM tumors was monitored with a 7T Bruker MRI at the MDACC Small Animal Imaging Facility (SAIF). Treatment started at the indicated times post-implantation and consisted of vehicle, 10% Captisol (Selleck) in 50 mM citrate buffer pH 5.0 (Teknova), 30 mg/kg MRTX1133 (WuXi AppTech) in vehicle, or 60 mg/kg MRTX1133 in vehicle (dose escalation) injected BID i.p. BI1347 (CDK8i, 10 mg/kg, MedChem Express) was administered QD by oral gavage and SB225002 (CXCR2i, 10 mg/kg, Selleck Chemicals) was administered QD i.p. in 10% Captisol in 50 mM citrate buffer pH 5.0. Anti-CTLA-4 (BioXcell, BE0131), anti-PD-1 (BioXcell, B0273) or the respective isotype (BioXcell, BE0087 for anti-CTLA-4; BioXcell, BE0089 for anti-PD-1) in PBS were administered three times per week i.p. once at a dose of 200 μg, followed by two doses of 100 μg. CD8 depletion experiments were previously described.^10^ Control arms for inhibitor experiments were administered vehicle and/or isotype antibody. PDXs were derived as previously described^55^ and implanted subcutaneously into 11 weeks old athymic nude mice. Tumor volumes were captured by serial caliper measurements (twice weekly) and treatment was initiated when tumors reached 150 to 250 mm^3^. Tumor volume (TV) was calculated as TV = (D × d2/2), where “D” is the larger and “d” is the smaller superficial visible diameter of the tumor mass. All measurements were documented as mm^3^. Treatment groups included 6 mice per group. To generate PDXs resistant to adagrasib, tumors were implanted and allowed to reach approximately 500 mm^3^ and dosed with adagrasib at 100 mg/kg QD by oral gavage or for 5 days on-treatment followed by 2 days off-treatment until tumors reached 1000 – 1500 mm^3^ in size. To generate MRTX1133-resistant PDXs, tumors were treated with MRTX1133 at 30 mg/kg BID by intraperitoneal injection for 5 days on-treatment followed by 2 days off-treatment until tumors reached 1000 – 1500 mm^3^. At the endpoint of the study, tumors were collected 4 hours after the last dose of treatment and fixed in 10% neutral buffered formalin overnight and then processed and embedded in paraffin. FFPE tissues were sectioned at 5 µm for histology analyses. All procedures were reviewed and approved by the MD Anderson Institutional Animal Care and Use Committee.

### Histological analysis

*5* μm-thick sections of formalin-fixed paraffin embedded (FFPE) tissues were stained with hematoxylin and eosin (H&E) with the S&T Infinity Staining and Leica Autostainer XL. Representative pictures were acquired with a Leica DM1000 LED microscope equipped with a DFC295 microscope camera (Leica) with LAS version 4.4 software (Leica). Five fields of view at 100x magnification per tissue were analyzed for histological scoring. Orthotopic models will have either PDAC or uninvolved tissue and were scored based on the percentage of the tissue that consisted of PDAC (1: no PDAC, 2: 0-25% PDAC, 3: 26-50% PDAC, 4: 51-75% PDAC, 5: 76-100% PDAC). Tissues from GEMMs were quantified based on the relative percentage of tissue with each indicated histological phenotype.

### Immunohistochemistry

For IHC staining, 5 μm formalin-fixed paraffin embedded (FFPE) tissues were processed as reported previously.^10^ Antigen retrieval was carried out by subjecting the sections to Tris-EDTA (TE) buffer (pH 9.0) for pERK, CD206, GFP, and MYC after deparaffinization and hydration. For CDK8 and CK19, antigen retrieval was performed with citrate buffer (pH 6.0). A hydrophobic barrier was then created around the tissue sections followed by the incubation of 3% H_2_O_2_ in PBS for 15 minutes to block endogenous peroxidase activity. The slides were blocked with 1.5% bovine serum albumin (Sigma Aldrich) for 30-60 minutes prior to incubation with anti-pERK (Cell Signaling 4376S, 1:250), CDK8 (Proteintech 22067-1-AP, 1:100), GFP (Abcam ab290, 1:400) CK19 (Abcam ab52625, 1:400) overnight at 4°C. For CD206 staining, blocking was performed in 4% cold water fish skin gelatin (CWFG, Aurion 14719NT) for 1 hour and staining with anti-CD206 (R&D Systems AF2535, 1:100) overnight at 4°C. For MYC staining, blocking was performed in 1% BSA for 1 hour and staining with anti-MYC (Cell Marque 395R-14, 1 μg/mL) overnight at 4°C. The slides were incubated with biotinylated anti-rabbit secondary antibody (Vector Laboratories BA-1000, 1:200-1:250) or biotinylated anti-goat secondary antibody (Jackson ImmunoResearch 705-066-147, 1:250) for 30 minutes in blocking buffer followed by incubation with ABC reagent (Vector Laboratories PK-6100) for 30 minutes. Subsequently, the slides were stained with DAB (Life Technologies), counterstained with hematoxylin, and cover slipped. Representative pictures were acquired with a Leica DM1000 LED microscope equipped with a DFC295 microscope camera with LAS version 4.4 software. At least three random fields for each tissue section were captured, and the staining per visual field (200X) was quantified with ImageJ (NIH, Bethesda, MD). Weighted IHC scoring was performed based on positive area (0: 0%, 1: 1-10%, 2: 11-50%, 3: 51-80%, 4: 81-100%) and intensity (0: negative, 1: weak, 2: moderate, 3: strong). Imaging and quantification were restricted to tumor regions for orthotopic models and in regions with PanIN/PDAC for KPCY models.

For the visualization of CK19, FAS, and CDK8 co-expression, tyramide signal amplification (TSA) multiplex staining was performed. 5 μm formalin-fixed paraffin embedded (FFPE) tumor tissues were deparaffinized and hydrated, antigen retrieval was performed in citrate buffer (pH 6.0) for 15 minutes at 98°C in a microwave. Antigen retrieval for CK19 on PDX samples was performed in TE buffer. Subsequently, a hydrophobic barrier was then created around the tissue sections before blocking with 4% CWFG for 1 hour at room temperature (RT), followed by the incubation with CK19 antibody (Abcam ab52625, 1:200) for 1 hour (PDX) or 3 hours (murine) at RT. Then the slides were incubated with rabbit on rodent HRP-Polymer (Biocare Medical) for 10 minutes (PDX) or 30 minutes (murine) at RT followed by staining with Opal 570 (Akoya Biosciences) for 10 minutes at RT. Then the slides were heated at 98°C in citrate buffer (pH 6.0) for 15 minutes in a microwave again followed by the blocking with 4% CWFG for 1 hour at RT, incubated with FAS antibody (PDX: Abcam ab133619, 1:1000; murine: Abcam ab271016, 1:100) overnight at 4°C (PDX) or for 3 hours at RT (murine), incubated with rabbit on rodent HRP-Polymer for 10 minutes (PDX) or 30 minutes (murine) at RT and Opal 690 (Akoya Biosciences) for 10 minutes at RT. Afterwards, the slides were heated at 98°C in citrate buffer (pH 6.0) for 15 minutes in a microwave followed by the blocking with 4% CWFG for 1 hour at RT, incubated with CDK8 (Proteintech 22067-1-AP, 1:100) for 1 hour (PDX) or 3 hours (murine) at RT, rabbit on rodent HRP-Polymer for 10 minutes (PDX) or 4plus Biotinylated Goat Anti-Rabbit IgG (Biocare Medical) for 30 minutes and 4plus Streptavidin HRP Label (Biocare Medical) for 30 minutes at RT (murine), stained with Opal 520 (Akoya Biosciences) for 10 minutes before the incubation with Hoechst 33342 (Invitrogen, 1:10,000) for 10 minutes and mounted with Vectashield (Vector Laboratories). A similar process was performed for CD8, Ki67, pERK, and CK19 staining, with antigen retrieval steps performed in TE buffer (pH 9.0) in a microwave at 98°C for 15 minutes and blocking in 1% BSA for 10 minutes, followed by either anti-CD8 (Cell Signaling 98941S, 1:800), anti-Ki67 (Thermo Fisher Scientific RM-9106-S, 1:400), pERK (Cell Signaling 4376, 1:500), or CK19 (Abcam ab52625, 1:400) incubation for 1 hour at room temperature or overnight at 4°C. Tissues were incubated with BioCare Rabbit-on-Rodent HRP polymer for 10 minutes at room temperature, and Opal 570 (for CD8 and pERK, 1:100) or Opal 520 (for Ki67 and CK19, 1:100) for 10 minutes at room temperature. Images were captured with a Zeiss LSM800 confocal microscope with a 20x objective and quantified by counting the number of positive cells in each visual field. For CK19 and pERK imaging, images were captured with 2x digital zoom (40x). Imaging and quantification were restricted to tumor regions for orthotopic models and in regions with PanIN/PDAC for KPCY models.

### PhenoCycler staining and analysis

Tissues were embedded in OCT and cryostat sectioned (5 μm) onto coverslips coated with 1X poly-L-lysine. For tissue retrieval, sample coverslips were placed over Drierite beads for 2 minutes before being immersed in acetone for 10 minutes and then placed in a humidity chamber for 2 minutes. Next, tissues were hydrated (two 2-minute incubations in Hydration Buffer, Akoya Biosciences), fixed [10 minutes, 1.6% paraformaldehyde (PFA) in hydration buffer solution], and rinsed in Hydration Buffer prior to staining.

For staining, tissues were first incubated for 20-30 minutes in Staining Buffer (Akoya Biosciences) before adding the CODEX antibody cocktail [CODEX antibodies (see **Table 3** for list of antibodies used) in CODEX Blocking Buffer, Akoya Biosciences] to coverslips and incubation in a humidity chamber at room temperature for 3 hours. Following the 3-hour antibody incubation, tissues were washed (2 times for 2 minutes) in Staining Buffer (Akoya Biosciences) before a secondary post-stain fixation step (10 minutes, 1.6% PFA in Storage Buffer, Akoya Biosciences). Next, tissues were washed (3 times with 1x PBS), incubated in ice-cold methanol for 5 minutes, and washed (3 times with 1x PBS) prior to a final fixation step in a humidity chamber [20 min at room temperature with Fixative Reagent (Akoya Biosciences) in 1x PBS]. Lastly, samples were washed 3 times in 1x PBS and placed in Storage Buffer (Akoya Biosciences) at 4°C.

**Table 3:** Antibodies used for PhenoCycler analysis.

| Antibody | Catalog # | Clone | Dilution |
| --- | --- | --- | --- |
| CD44 | 4250002 | IM7 | 1:375 |
| CD3 | 4550109 | 17A2 | 1:200 |
| CD4 | 4250016 | RM4-5 | 1:200 |
| CD5 | 4250007 | 53-7.3 | 1:200 |
| CD8a | 4250017 | 53-6.7 | 1:200 |
| CD11b | 4150015 | M1/70 | 1:200 |
| CD11c | 4550108 | N418 | 1:200 |
| CD19 | 4250014 | 6D5 | 1:200 |
| Ly6G | 4550110 | 1A8 | 1:200 |
| TCRβ | 4550101 | H57-597 | 1:200 |
| MHCII | 4250003 | M5/114.15.2 | 1:200 |

Stained coverslips were run on the PhenoCycler System (Akoya Biosciences) to obtain stitched 20X images (Keyence, BZ-X800) of barcoded reporters. Images were analyzed using QuPath. Classifiers for markers were trained by researchers blinded to sample group and applied to images to quantify individual staining as well as co-expression of markers. Raw counts were then normalized to tissue area (mm^2^).

### scRNA-seq sample preparation

Tissue digestion and scRNA-seq preparation was performed as previously described.^10^ Briefly, 2-3 pooled tumors per condition were minced with a razor and digested with 4 mg/mL collagenase IV and 4 mg/mL dispase in RPMI with 1% PSA for 50 minutes at 37°C with shaking, followed by consecutive filtering through 70 μm and 40 μm filters. Samples were washed with RPMI with 10% FBS and 1% PSA, centrifuged at 500g for 5 minutes, washed with FACS buffer (PBS with 2% FBS), centrifuged at 500g for 5 minutes, and stained with Fixable Viability Dye eFluor™ 780 (eBioscience, 1:1,000) on ice for 10 minutes. Samples were washed twice with washed with FACS buffer, centrifuged at 500g for 5 minutes, and sorted with a BD FACSMelody. Cell counting was performed with Trypan Blue (Sigma Aldrich) exclusion with a Countess 3 (Invitrogen). 5,000 to 10,000 cells were loaded into the 10x Chromium Controller to generate Single Cell 3’ Gene Expression dual indexed libraries (10x Genomics). The creation of Gel Bead-In-Emulsions (GEM) and subsequent cell barcoding, GEM-RT cleanup, cDNA amplification, and library assembly were performed with the Chromium Next GEM Single Cell 3’ Reagents Kits v3.1 (10x Genomics) according to manufacturer’s instructions. Paired-end, dual indexing sequencing was carried out using the Illumina NovaSeq6000.

### scRNA-seq data processing and analysis

scRNA-seq data processing and analysis was performed as previously described.^10^ Gene imputation was conducted by Markov affinity-based graph imputation of cells (MAGIC).^56^ Seurat employs a global-scaling normalization method “LogNormalize” that normalizes the feature expression measurements for each cell by the total expression, multiplies this by a scale factor (10,000 by default), and log-transforms the result. To have a comparable number of cells for all samples included in the same comparison analysis, we downsampled the HY19636 MRTX1133+CDK8i+αCTLA-4 sample by randomly selecting 1400 cells from the original dataset using the R function “sample”. We also downsampled the orthotopic KPC689 resistant MRTX1133 sample to 3300 cells to have a similar number of cells to KPC689 parental. GSEA was performed with GSEA 4.2.2^57,58^ based on differentially expressed genes for each cluster and pathways with an FDR q-value < 0.1 reported.

### Visium spatial transcriptomics analysis

H&E stained 5 μm thick FFPE tissues were scanned with a Zeiss Axio Scan.Z1 followed by processing with the Visium CytAssist Spatial Gene Expression for FFPE, Mouse Transcriptome kit (10x Genomics) according to manufacturer’s instructions. Prepared libraries were sequenced with a NextSeq500 (Illumina), NextSeq2000 (Illumina), or NovaSeq6000 (Illumina) according to manufacturer’s recommendations.

Raw FASTQ files were mapped to the mm10 reference genome and spatially projected using Space Ranger 2.0.1 (10x Genomics). Cell2location (version 0.1.3) (PMID: 35027729) was employed to create spatial maps of cell types by integrating Visium spatial data and scRNA-seq references of the matched samples. Cell2location decomposes multi-cell spatial transcriptomics data by estimating the abundance of reference cell types at specific spatial coordinates using a principled Bayesian model. Our analysis includes multiple steps. First, a high-quality reference of cell annotations was constructed using the scRNA-seq data of matched mouse samples and fed into cell2location as input. Second, cell2location performed gene selection and defined expression signatures for each cell type. Subsequently, the default negative binomial regression model in cell2location was utilized to estimate the reference cell-type signatures for determining cell-type locations. For the analysis of each Visium section, the default parameters were applied, with the exception of the following settings: ‘max_spochs’: 50000; ‘N_cells_per_location’: 5; ‘detection_alpha’: 20. To visually represent the contribution of each cell type to each spot, the 5% percentile of the posterior distribution of mRNA counts was employed. The gene expressions of each cell type at each spot were also estimated using cell2location with the adopted approach from robust cell type decomposition (RCTD) (PMID: 33603203). The estimated gene expressions of each cell type were exported and the spots stratified based on the mean *Cdk8* expressions in ductal/cancer cells by excluding the spots with zeros. The expression and cell proportion matrices were concatenated for each spot and the combined matrix input back to cell2location for visualization.

### In vitro MRTX1133 treatment

For in vitro experiments with MRTX1133 treatment, 5×10^5^-1×10^6^ cells were seeded in one well of a 6 well plate. 24 hours later, cells were treated with vehicle (DMSO) or the indicated concentrations of MRTX1133 in DMSO. RNA was collected at 6- or 24-hours post-treatment according to the RNeasy kit protocol (Qiagen). cDNA synthesis was performed with the High-Capacity cDNA Reverse Transcription kit (Applied Biosystems) and qPCR performed with Power SYBR Green PCR Master Mix and the primers listed in **Table 4** and analyzed with the Applied Biosystems Quant Studio 7 Flex Real-Time PCR System (Applied Biosystems). Gene expression is calculated using the 2^−ΔΔCT^ method with *Gadph* or *18s* as a housekeeping gene and normalized to vehicle or shNT as indicated in the figure legends.

**Table 4:**
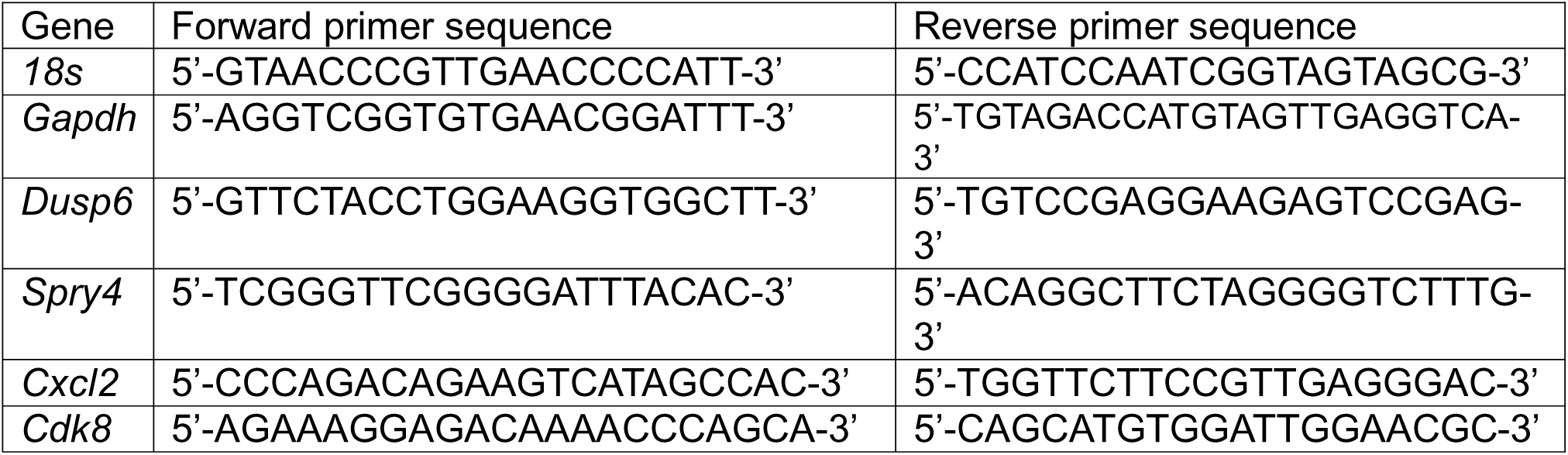
qPCR primer sequences.

Protein was collected at 3 hours post-treatment in urea lysis buffer (8 M urea, 2.5% SDS) with cOmplete mini EDTA-free protease inhibitor cocktail (Roche) and quantified by Pierce BCA assay according to manufacturer’s instructions (Thermo Scientific). 2 μg/μL of protein lysate was loaded and samples prepared according to the manufacturer’s instructions with the anti-rabbit detection module (Bio-Techne) and the antibodies listed in **Table 5**. Analysis was performed with the Wes instrument (Bio-Techne) and Compass for SW software (Bio-Techne). Quantification of pERK was performed in the Compass for SW software, where pERK peaks were normalized to vinculin peaks. Further normalization to the DMSO treated sample for each cell line was performed.

**Table 5:** Simple Western antibodies.

| Antibody | Manufacturer | Catalog # | Dilution |
| --- | --- | --- | --- |
| Ras, mutant G12D specific | Cell Signaling Technology | 14429 | 1:10 |
| Phospho-p44/42 MAPK (Erk1/2)<br>(Thr202/Tyr204) | Cell Signaling Technology | 4370 | 1:10 |
| p44/42 MAPK (Erk1/2) | Cell Signaling Technology | 9102 | 1:10 |
| Vinculin | Abcam | ab129002 | 1:50 |

In other experiments, 5×10^5^-1×10^6^ cells were seeded per well of a 6 well plate and 24 hours later, treated with MRTX1133 for 3 hours, 6 hours, or 24 hours. Cells were lysed in urea lysis buffer (8 M urea, 2.5% SDS) with cOmplete mini EDTA-free protease inhibitor cocktail and quantified by Pierce BCA assay according to manufacturer’s instructions. 20 μg of protein was loaded per sample onto Bolt 4-12% Bis-Tris Plus gels (Invitrogen) and transferred using the Trans-Blot Turbo RTA Mini PVDF Transfer Kit (Bio-Rad). Membranes were blocked in 5% BSA in TBS with 0.1% Tween-20 (TBS-T) and incubated in the primary antibodies listed in **Table 6** overnight at 4°C. Membranes were washed with TBS-T, followed by incubation in secondary antibody (**Table 6**) for 1 hour at room temperature. Membranes were washed with TBS-T and developed with West-Q Pico ECL Solution (GenDepot). Following development of each antibody, membranes were stripped with 1.5% glycine (pH 2.2), 0.1% SDS, and 1% Tween-20, blocked, and incubated with primary and secondary antibody as described above. Membranes were first probed for RAS^G12D^, followed by pERK, ERK, and vinculin.

**Table 6:** Western blot antibodies.

| Antibody | Manufacturer | Catalog # | Dilution |
| --- | --- | --- | --- |
| Phospho-p44/42 MAPK (Erk1/2) (Thr202/Tyr204) | Cell Signaling Technology | 4370 | 1:1,000 |
| p44/42 MAPK (Erk1/2) | Cell Signaling Technology | 9102 | 1:1,000 |
| Ras, mutant G12D specific | Cell Signaling Technology | 14429 | 1:1,000 |
| Vinculin | Abcam | ab129002 | 1:1,000 |
| Anti-rabbit IgG, HRP | Abcam | ab16284 | 1:1,000 |

For proliferation analysis, 5×10^4^ cells/mL were seeded in 50 μL of DMEM with 2.5% FBS, 1% PSA, 1 mM sodium pyruvate (Gibco) in 96-well white bottom plates (Corning). 24 hours later, media was removed, and cells treated with varying concentrations of MRTX1133 in DMEM with 2.5% FBS, 1% PSA, 1 mM sodium pyruvate (50 μL total volume, with 8 μg/mL puromycin added for shCdk8 experiments). After 3 days, plates were equilibrated to room temperature for 30 minutes followed by addition of 50 μL of CellTiter-Glo Reagent according to manufacturer’s instructions (Promega). Luminescent signal was measured with a FLUOstar Omega plate reader (BMG LABTECH), the signal in media only wells subtracted, followed by normalization to wells treated with DMSO.

### Digital droplet PCR and whole exome sequencing

DNA was isolated from cells or snap frozen pancreas/tumor tissue using the DNeasy Blood and Tissue Kit (Qiagen) and quantified with the Qubit dsDNA HS Assay Kit (Invitrogen) according to the manufacturer’s instructions. 10 ng of genomic DNA (gDNA) was mixed with 1x QuantStudio 3D Digital PCR Master Mix v2 and 1x *KRAS^G12D^*TaqMan dPCR Liquid Biopsy Assay (Applied Biosystems A44177, Hs000000051), loaded onto QuantStudio 3D Digital PCR 20K chips (Applied Biosystems), and analyzed with the QuantStudio 3D Digital Real-Time PCR System (Applied Biosystems) and QuantStudio 3D Analysis Suite software (Applied Biosystems). A no template control was used to establish thresholds for the assay.

Mouse Exome panel (Twist Biosciences) was used for whole-exome sequencing according to manufacturer’s instructions. Samples were sequenced with the NovaSeq6000. For data analysis, quality control was conducted for the FASTQ files using FastQC (version 0.11.9)^59^ and adapter sequences were removed by Cutadapt (version 4.1)^60^. The processed FASTQ files were then mapped to the reference genome (mm10) using bwa mem (version 0.7.17, arXiv:1303.3997v2). The PCR duplicate reads were marked by Picard (version 2.27.4)^61^ and base recalibration was performed using GATK Base Quality Score Recalibration (BQSR) (GATK 4.1.7.0)^62^. Copy number variation analysis was performed by CNVkit (version 0.9.9)^63^ for each tumor and cell line sample using the matched tail gDNA sample and parental cell line as reference. The plots related to CNV analysis were generated by the R package ggplot2 (version 3.5.0).

### Quantification and statistical analysis

Graphical representation of the data and statistical tests were performed using GraphPad Prism 10 and reported in the respective figure legends. Shapiro-Wilk test was used to assess the normality of distribution of samples. Two-sided unpaired t-test (for samples with normal distribution and equal variances), unpaired t-test with Welch’s correction (for normally distributed samples with unequal variances), or two-sided Mann-Whitney test (for samples with non-normal distribution) were performed for comparison of two samples. For comparison of more than 2 groups, one-way analysis of variance (ANOVA) with Tukey’s or Dunnett’s multiple comparisons test (for samples with normal distribution), Brown-Forsythe and Welch ANOVA with Dunnett’s T3 multiple comparisons test (for samples with unequal variances) and Kruskal-Wallis with Dunn’s multiple comparisons test (for samples with non-normal distribution) were used. Two-way ANOVA with Sidak’s multiple comparisons test was used. Log-rank test or Gehan-Breslow-Wilcoxon test were used to compare Kaplan-Meier survival curves. P-values are reported as * P < 0.05, ** P < 0.01, *** P < 0.001, **** P < 0.0001, *ns*: not significant. Exact p-values are reported where indicated.

## Data availability

scRNA-seq data are deposited at GEO (GSE269679) and spatial transcriptomics data are deposited at GEO (GSE269680). WES data was deposited at SRA (Bioproject <u>PRJNA1124144).</u>

## Acknowledgements

This work was supported by Break *Through* Cancer and the Lustgarten Foundation for Pancreatic Cancer Research. KKM and KMM were supported by Ergon Foundation Post-Doctoral Trainee Fellowships. RK is a Distinguished University Chair supported by Sid W. Richardson Foundation. The Kalluri laboratory PDAC related research is supported by NCI P01 CA117969. AM is also supported by the Sheikh Khalifa bin Zayed Foundation and NCI U54CA274371. Other support includes the Small Animal Imaging Facility and Advanced Technology Genomics Core at MD Anderson Cancer Center supported by NCI P30CA16672. We thank Haoqiang Ying for providing the HY19636 cell line and Anais Comptdaer for assistance with in vitro experiments.

## Author contributions

KMM, KKM, MLK, HY, and DP designed, performed, and supervised in vivo studies. AMS, SJM, SIP, BAMD, HS, and AAB performed in vivo experiments and analyzed data. KMM, KKM, AMS, BAMD, XZ, LKS, MRC, SIP, and SK performed histological analyses. BL performed scRNA-seq, spatial transcriptomics, and whole-exome sequencing analyses. KMM, PJK, FP, and VB performed sample preparation for spatial transcriptomics analysis. KMM, KAA, YB, ANCA, KEB, and PTT performed in vitro experiments. KMM, KAA, and NS performed sample preparation and library preparation for scRNA-seq. KMM, PJK, and NS performed sequencing. KMM, KKM, AM, TPH and RK conceived and designed the project, and wrote or edited the manuscript. KMM, KKM, and BL generated figures.

