## Supplemental Figures for "CDK8 remodels the tumor microenvironment to resist the therapeutic efficacy of targeted KRAS^G12D^ inhibition in pancreatic ductal adenocarcinoma"

#### Supplemental Figure Legends

#### Supplemental Figure 1

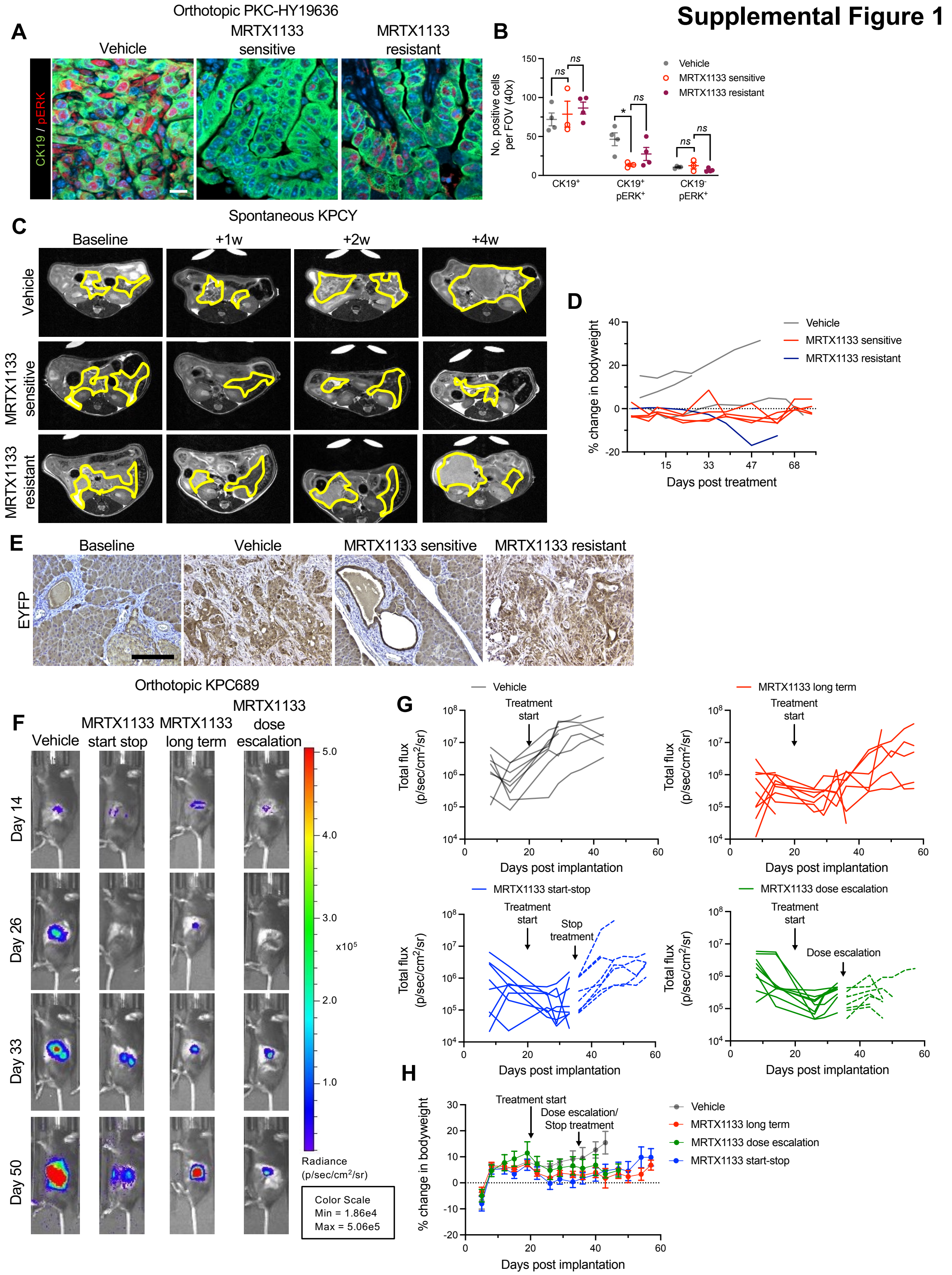


#### Supplemental Figure 1: Anti-PDAC responses of spontaneous and orthotopic KPC

**models to MRTX1133.** (**A-B**) Representative immunostaining images (**A**) and quantification of $CK19^{+}$, $CK19^{+}pERK^{+}$, and $CK19^{-}pERK^{+}$ cells in orthotopic PKC-HY19636 tumors (**B**). Vehicle, n=4; MRTX1133 sensitive, n=3; MRTX1133 resistant, n=4. Scale bar, 20 μm. (**C**) MRI of spontaneous KPCY models. Yellow lines demarcate tumor regions. (**D**) Bodyweight of KPCY mice during course of treatment. Vehicle, n=3; MRTX1133 sensitive, n=4; MRTX1133 resistant, n=1. (**E**) Representative images of EYFP lineage tracer stains of spontaneous KPCY tumors. Scale bar, 100 μm. (**F-G**) Representative bioluminescence images (**F**) and individual bioluminescence curves (**G**) of KPC689 tumors. Vehicle: n=8, MRTX1133 long term: n=9, MRTX1133 start-stop: n=8, MRTX1133 dose escalation: n=9. (**H**) Bodyweight of mice bearing orthotopic KPC689 tumors post tumor-implantation. Vehicle: n=8, MRTX1133 long term: n=9, MRTX1133 start-stop: n=8, MRTX1133 dose escalation: n=9. Data are presented as mean ± s.e.m. except in (D) and (G) where individual mice are plotted. One-way ANOVA with Sidak's multiple comparisons test performed in (B). * $P < 0.05$, *ns*: not significant.

#### Supplemental Figure 2

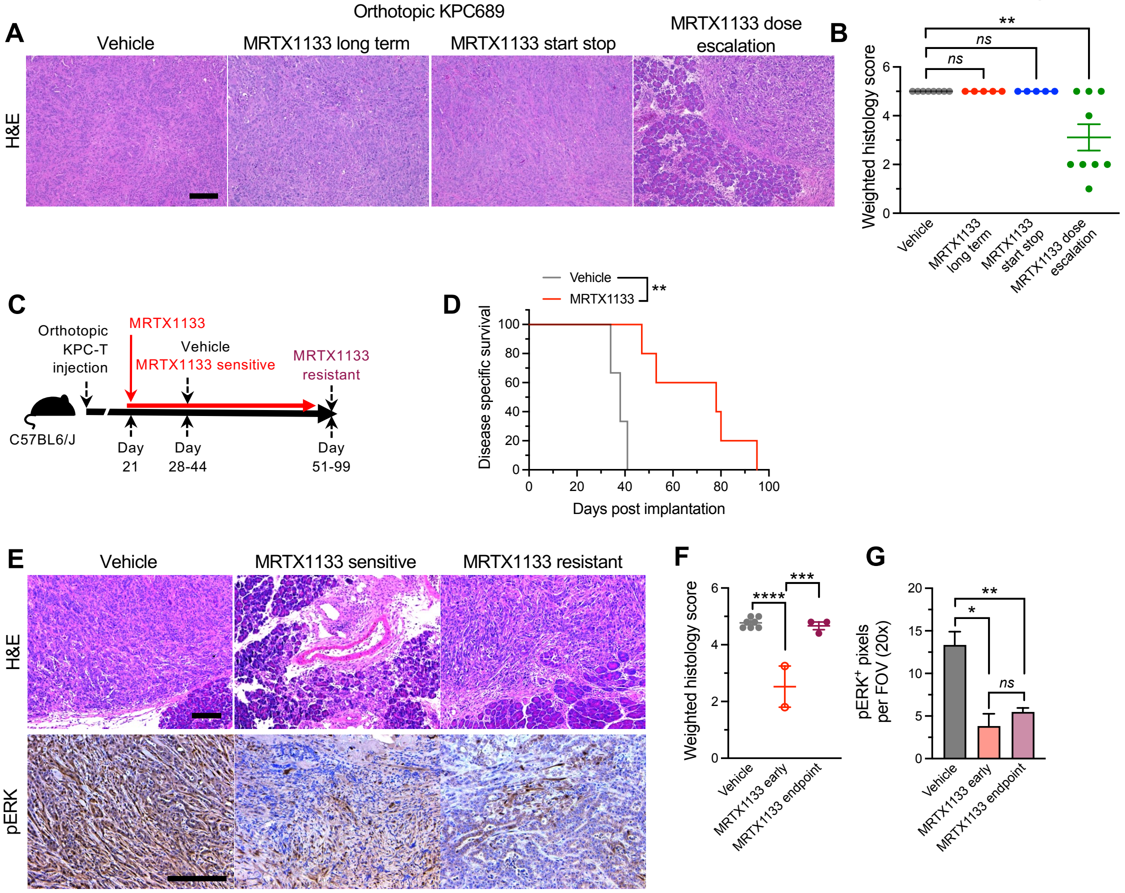


**Supplemental Figure 2: Response of orthotopic tumors to MRTX1133.** (**A-B**) Representative H&E images (**A**) and quantification (**B**) of orthotopic KPC689 tumors. Vehicle: n=8, MRTX1133 long term: n=5, MRTX1133 start-stop: n=5, MRTX1133 dose escalation: n=9. (**C**) Schematic of orthotopic KPC-T tumors treated with MRTX1133. (**D**) Survival curve of orthotopic KPC-T mice. Vehicle, n=3, MRTX1133, n=5. Note: Same mice from Fig. 5E are repeated here to allow for direct comparisons. (**E-G**) Representative H&E images (**E**) and quantification (**F**) of orthotopic KPC-T tumors. Vehicle, n=7; MRTX1133 sensitive, n=2; MRTX1133 resistant, n=3. Representative pERK staining (**E**) and quantification (**G**) of orthotopic KPC-T tumors. Vehicle, n=7, MRTX1133 sensitive, n=2, MRTX1133 resistant, n=5. Kruskal-Wallis ANOVA with Dunn's multiple comparison test performed in (B). Log-rank test performed in (D). One-way ANOVA with Tukey's multiple comparisons test performed in (F) and (G). Scale bars, 100 μm. Data are presented as mean ± s.e.m.* $P<0.05$, ** $P < 0.01$, *** $P < 0.001$, *ns*: not significant.

### Supplemental Figure 3

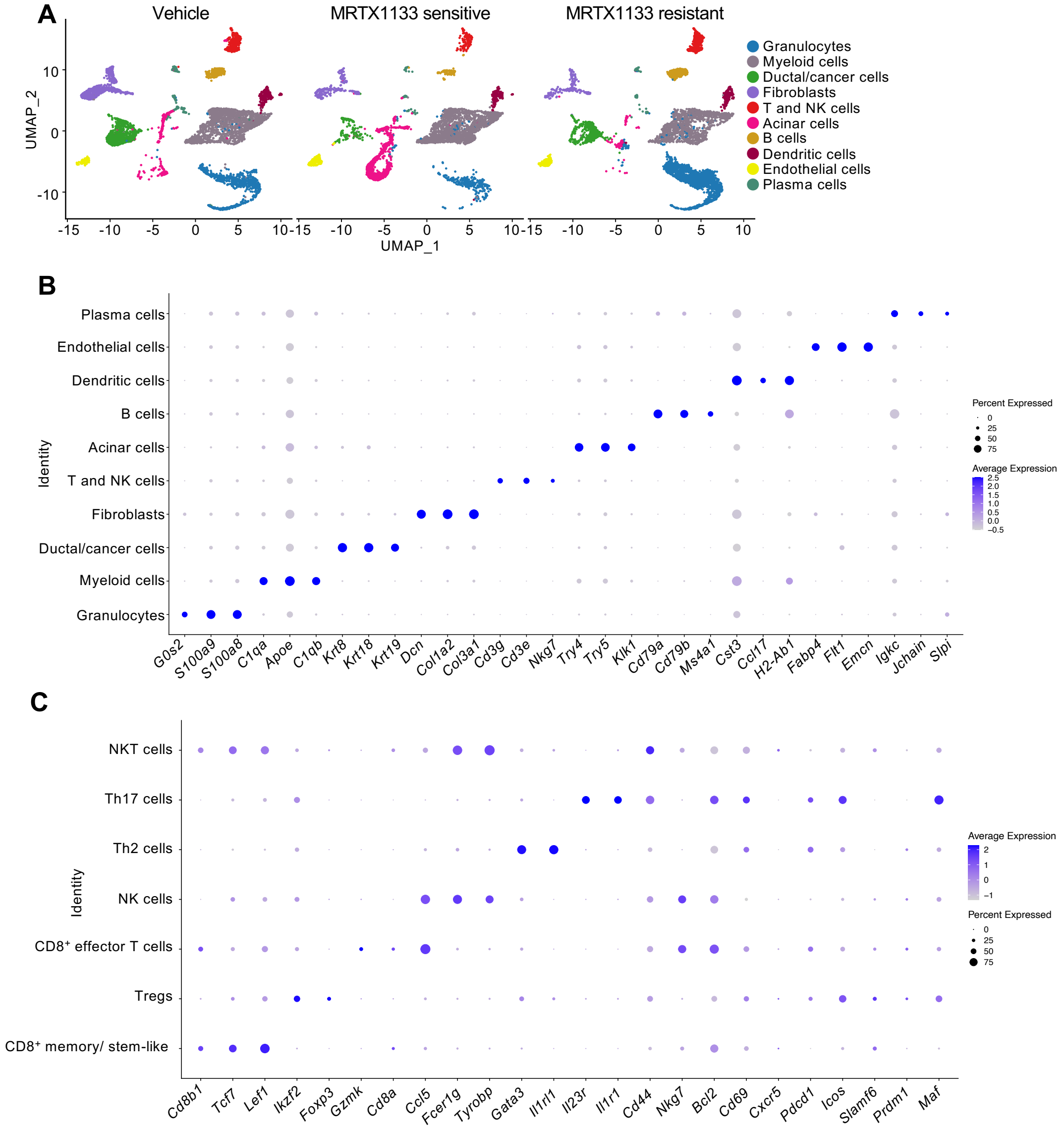


**Supplemental Figure 3: Cell type annotations based on scRNA-seq of orthotopic PKC-HY19636 tumors.** (**A-C**) UMAPs (**A**) of orthotopic PKC-HY19636 tumors determined by scRNA-seq. Dot plot of genes used to define general cell types (**B**), T and NK cell subtypes (**C**) of orthotopic PKC-HY19636 tumors determined by scRNA-seq.

#### Supplemental Figure 4

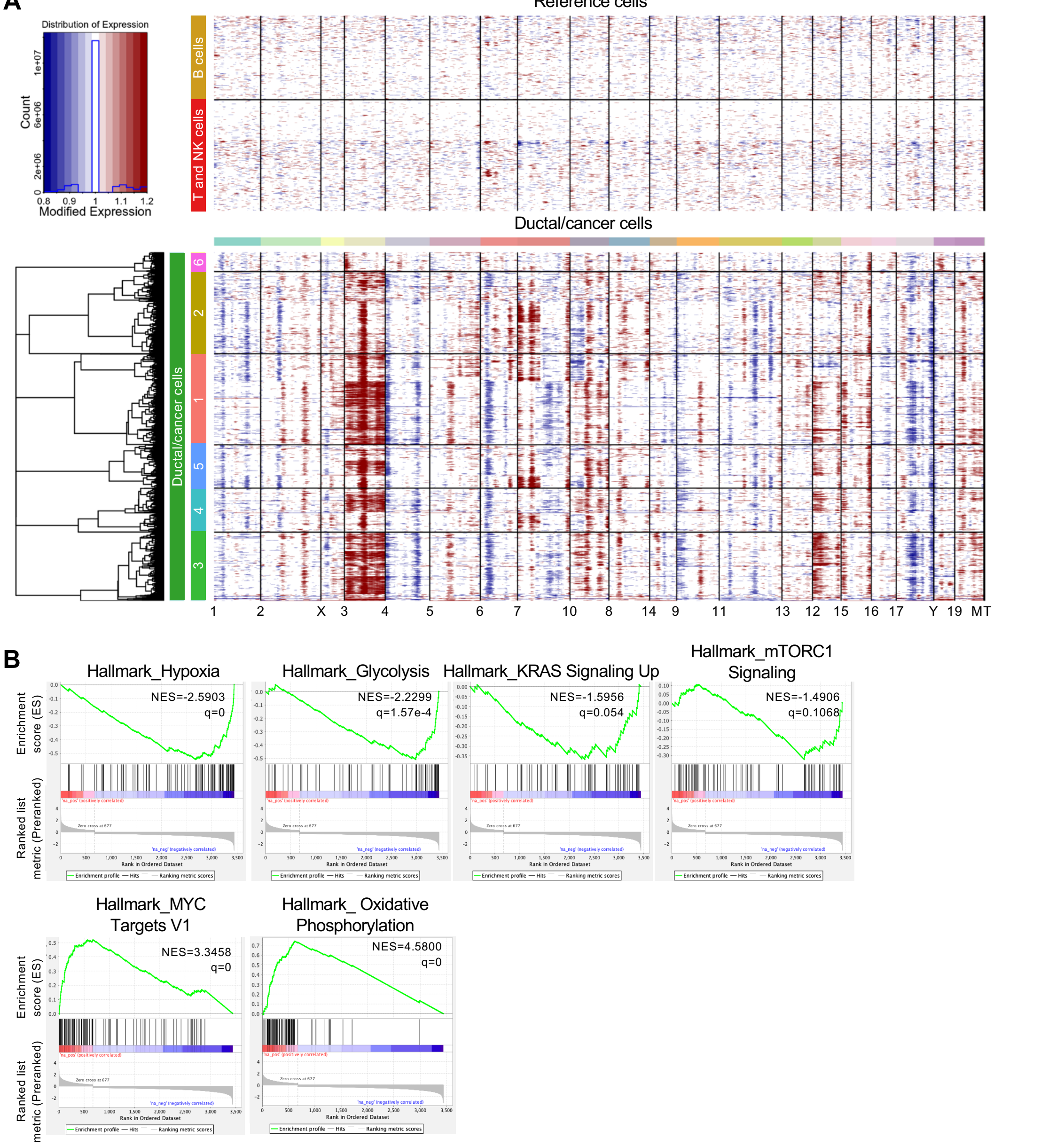


**Supplemental Figure 4: InferCNV analysis and enriched pathways of ductal/cancer cells based on scRNA-seq of orthotopic PKC-HY19636 tumors.** (**A**) InferCNV analysis of the ductal/cancer cell cluster of PKC-HY19636 tumors defined in Fig. 3A-B and Supplemental Fig. 2A-B. The T cell and B cell clusters were used as a reference.

Ductal/cancer cell clusters match the clusters defined in Fig. 3B. (**B**) GSEA of pathways enriched in ductal/cancer cells of MRTX1133 resistant orthotopic PKC-HY19636 tumors.

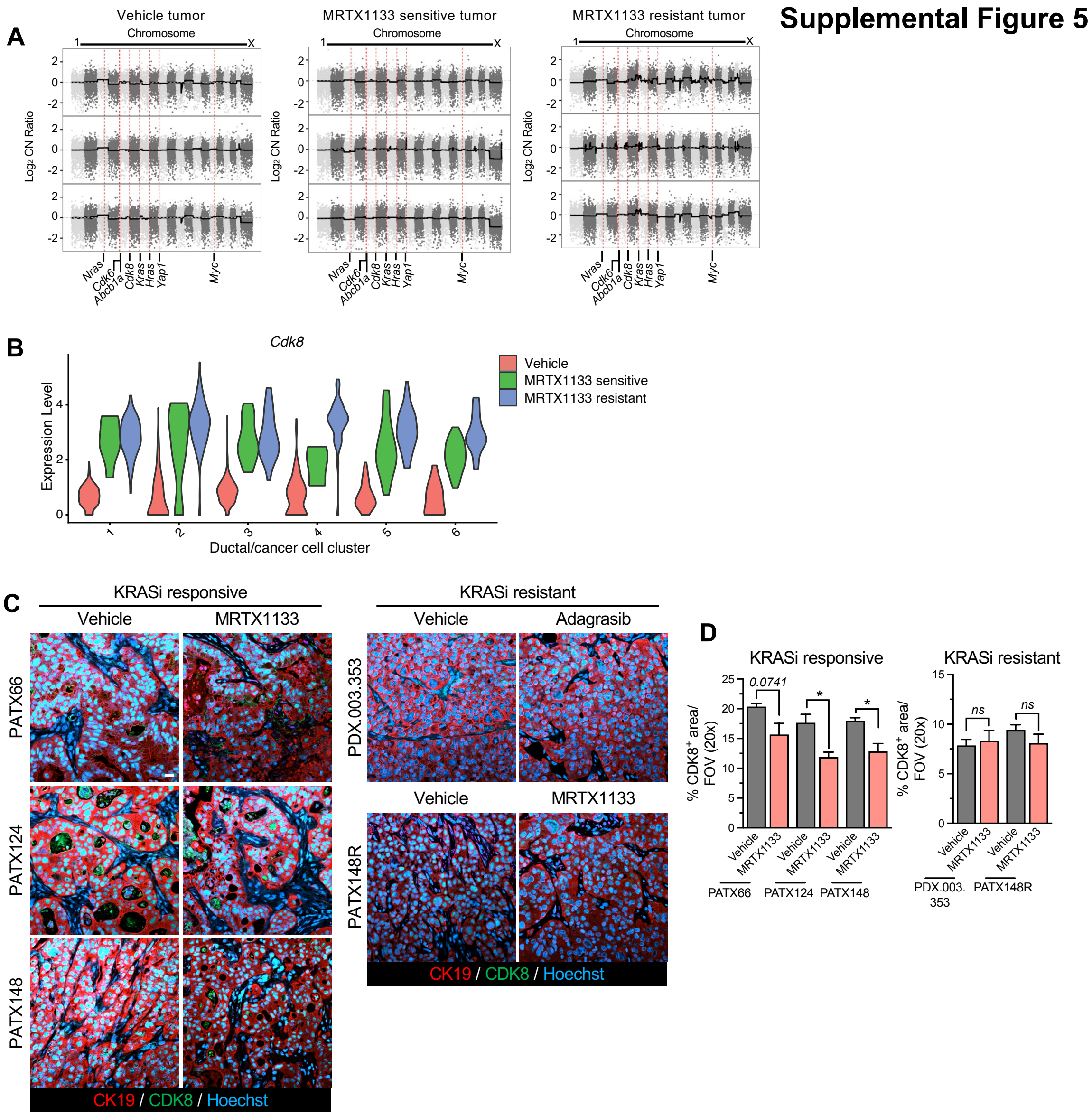


**Supplemental Figure 5: CDK8 is upregulated in PDAC resistant to MRTX1133.** (**A**) Genome-wide copy number plots of vehicle, MRTX1133 sensitive, and MRTX1133 resistant orthotopic PKC-HY19636 tumors with tail gDNA used as a reference. n=3 tumors per group. (**B**) Violin plot of *Cdk8* expression in ductal/cancer cell clusters of vehicle, MRTX1133 sensitive, and MRTX1133 resistant orthotopic PKC-HY19636 tumors analyzed by scRNA-seq. **(C-D)** Representative immunofluorescent images (**C**) and quantification (**D**) of KRASi responsive and KRASi resistant PDXs treated with vehicle, MRTX1133, or adagrasib. Scale bar, 20 µm. Unpaired t-test performed in (D). * $P < 0.05$, *ns*: not significant, or exact p-value reported.

#### Supplemental Figure 6

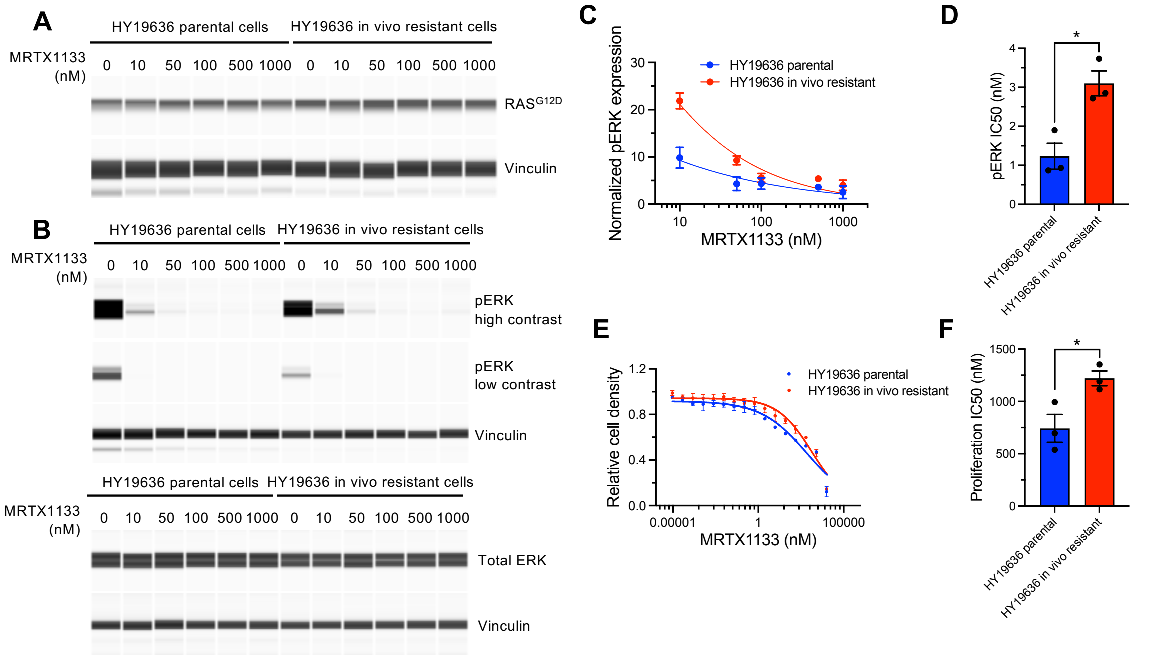


**Supplemental Figure 6: MRTX1133 resistant cells isolated from tumors demonstrate increased pERK and proliferation in response to MRTX1133 in vitro.** (**A**) Simple Western immunoassay for $RAS^{G12D}$ and vinculin of HY19636 parental and HY19636 in vivo resistant cells treated with the indicated concentrations of MRTX1133 for 3 hours. (**B**) Simple Western immunoassay for pERK, total ERK, and vinculin of HY19636 parental and HY19636 in vivo resistant cells treated with the indicated concentrations of MRTX1133 for 3 hours. A low contrast and high contrast image of the same gel are shown to better visualize samples with low pERK expression. (**C-D**) pERK expression normalized to vinculin and the vehicle treated (0 nM) control for each cell line

(**C**) and quantification of pERK IC50 (**D**). n=3 independent experiments. (**E-F**) Relative cell density of HY19636 parental and HY19636 in vivo resistant cells treated with the indicated concentrations of MRTX1133 for 3 days (**E**) and quantification of proliferation IC50 (**F**). n=3 independent experiments. Unpaired two-tailed t-test performed in (D) and (F). Data are presented as mean ± s.e.m. * $P < 0.05$.

#### Supplemental Figure 7

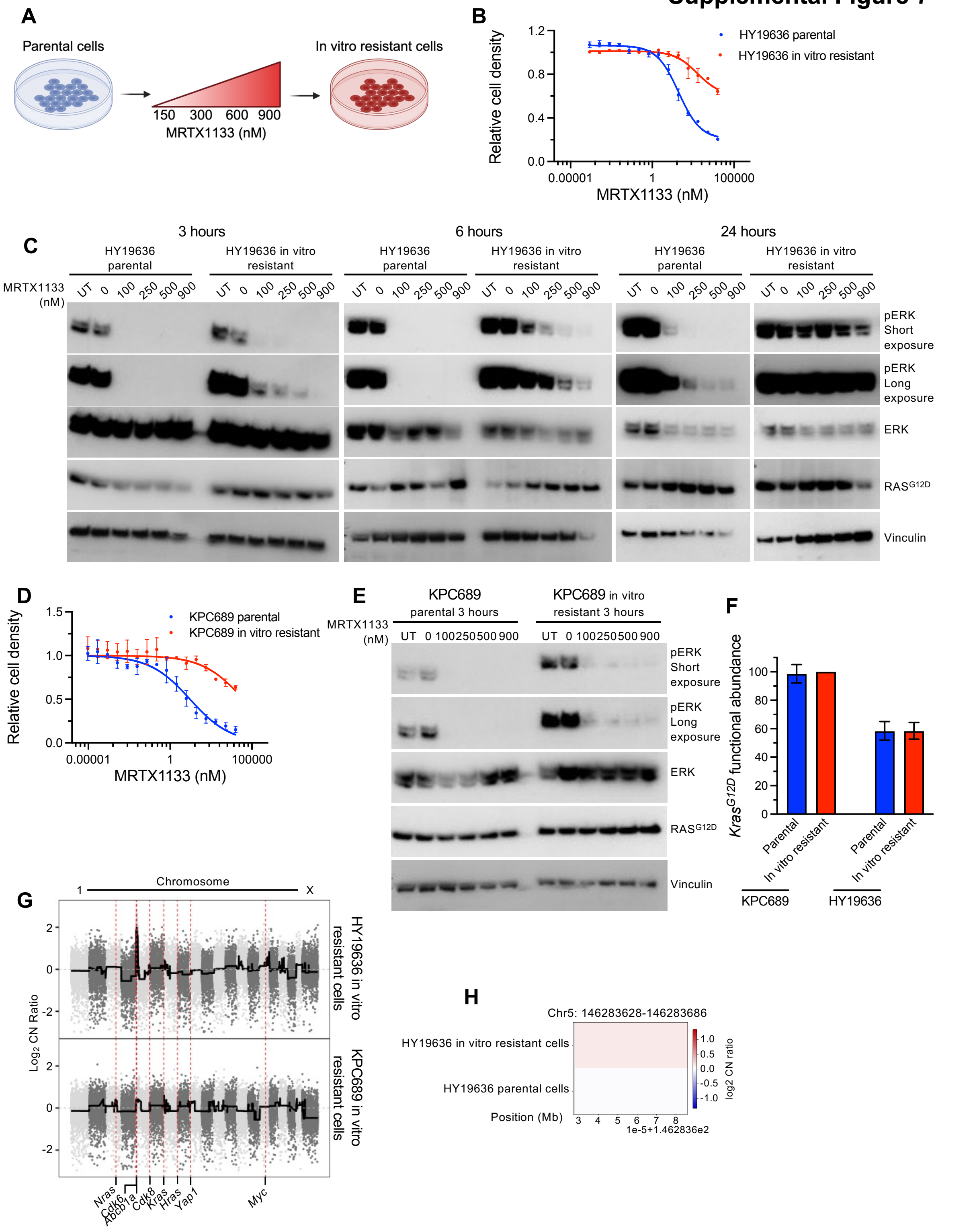


**Supplemental Figure 7: Characterization of in vitro MRTX1133 resistant cells.** (A)

Schematic of establishment of in vitro resistant cell lines, which were generated by exposing cells to increasing concentrations of MRTX1133 over the course of 2-3 months. (**B**) Relative cell density of HY19636 parental and HY19636 in vitro resistant cells treated with the indicated concentrations of MRTX1133 for 3 days. (**C**) Representative western blots of pERK, ERK, $RAS^{G12D}$, and vinculin of HY19636 parental and HY19636 in vitro resistant cells treated with the indicated concentrations of MRTX1133 for 3 hours, 6 hours, or 24 hours. (**D**) Relative cell density of KPC689 parental and KPC689 in vitro resistant cells treated with the indicated concentrations of MRTX1133 for 3 days. (**E**) Representative western blots of pERK, ERK, $RAS^{G12D}$, and vinculin of KPC689 parental and KPC689 in vitro resistant cells treated with the indicated concentrations of MRTX1133 for 3 hours. (**F**) ddPCR for $\textit{Kras}^{G12D}$ functional abundance in KPC689 and HY19636 parental and in vitro resistant cells. (**G**) Genome-wide copy number plots of HY19636 and KPC689 in vitro resistant with parental cell lines used as a reference. (**H**) WES data of the *Cdk8* gene of HY19636 parental and HY19636 in vitro resistant cells. Data are presented as mean ± s.d.

**Supplemental Figure 8**

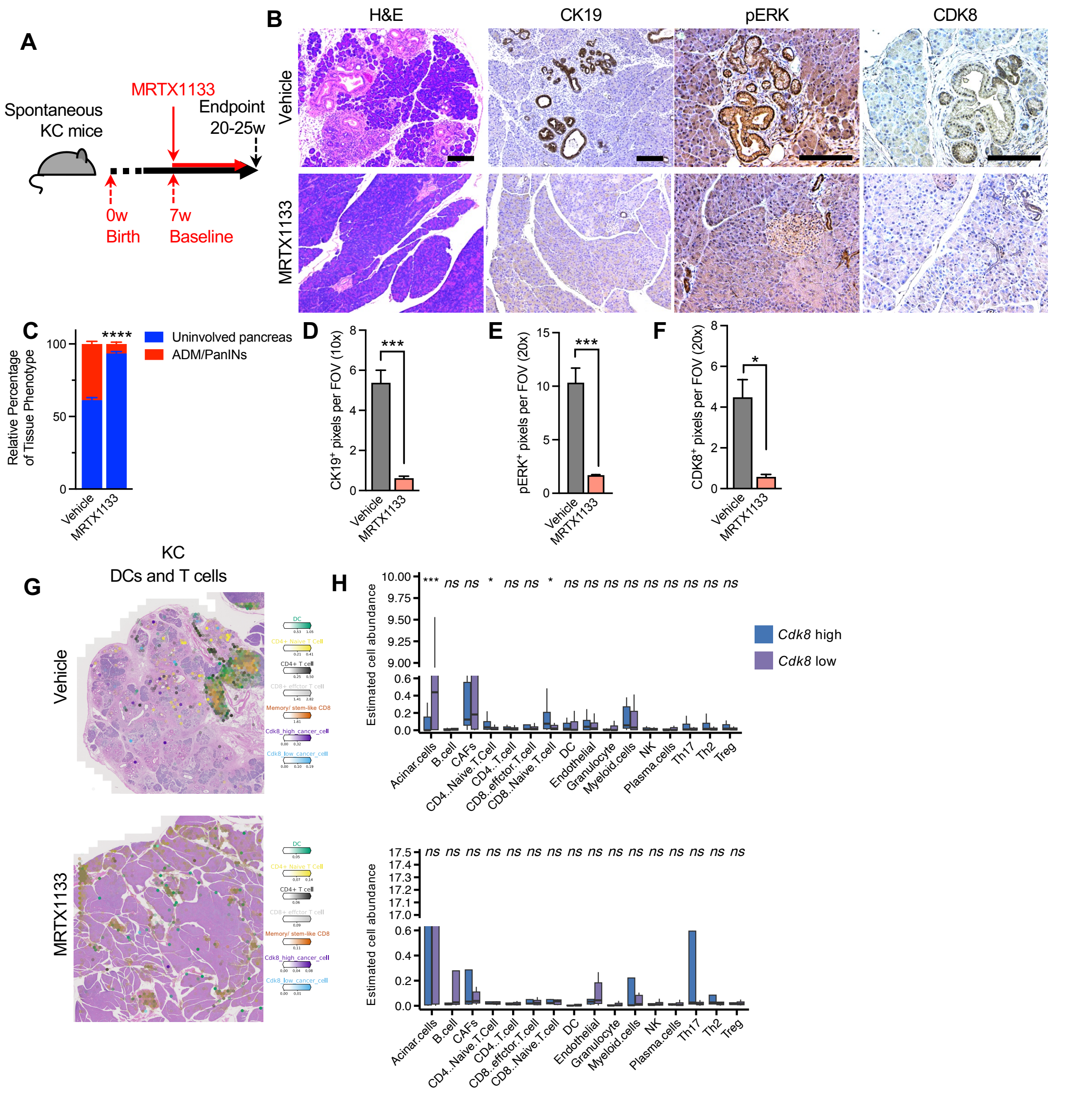


**Supplemental Figure 8: Response of KC pancreata to MRTX1133**. (**A**) Schematic of spontaneous KC mice treated with MRTX1133. (**B-F**) Representative H&E, CK19, pERK, and CDK8 images (**B**) and quantification (**C-F**) of KC pancreata. C: Vehicle, n=2; MRTX1133, n=4. D: Vehicle, n=2; MRTX1133, n=4. E: Vehicle, n=2; MRTX1133, n=4. F: Vehicle, n=5; MRTX1133, n=5. (**G-H**) Spatial transcriptomics representative images (**G**) and quantification (**H**) of KC pancreata. Spots were stratified by *Cdk8* high and low areas.

Scale bar, 100 µm. Two-sided Mann-Whitney test was performed in (H). Two-way ANOVA with Sidak's multiple comparisons test performed in (C). Unpaired t-test performed in (D) and (E). Unpaired t-test with Welch's correction performed in (F). Data are presented as mean ± s.e.m. except for in (H) where it is presented as median and range. * P < 0.05, *** P < 0.001, *ns*: not significant.

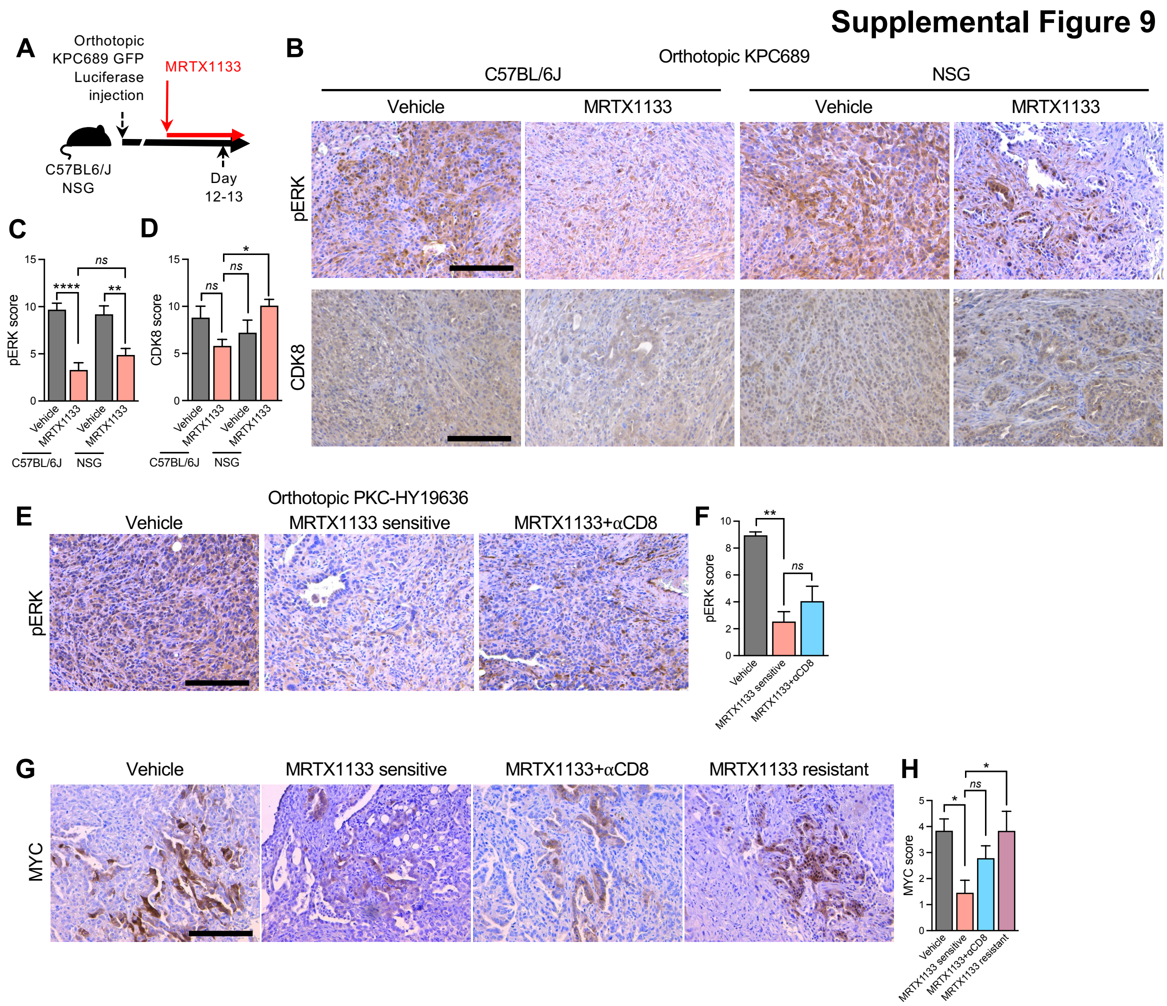


**Supplemental Figure 9: T cells control the emergence of CDK8⁺ resistant cancer cells**. (**A**) Schematic representation of orthotopic KPC689 GFP-luciferase tumors in C57BL6/J and NSG mice treated with MRTX1133. (**B**) Representative pERK and CDK8

staining at 12-13 days post-treatment. (**C-D**) Quantification of pERK (**C**) and CDK8 (**D**) staining. C57BL6/J MRTX1133, n=4 mice; n=5 mice per group for all other groups. (**E**) Schematic representation of orthotopic PKC-HY19636 tumors treated with MRTX1133 for 5-9 days. (**F-G**) Representative pERK staining (**F**) and quantification (**G**) of orthotopic PKC-HY19636 tumors. (**H-I**) Representative CDK8 staining (**H**) and quantification (**I**) of orthotopic PKC-HY19636 tumors. Vehicle, n=7 mice; MRTX1133 sensitive, n=4 mice; MRTX1133+αCD8, n=4 mice. Scale bars, 100 μm. One-way ANOVA with Dunnett's multiple comparisons test performed in (C), (D), (G), and (I). * $P < 0.05$, ** $P < 0.01$, *** $P < 0.001$, **** $P < 0.0001$, *ns*: not significant.

#### Supplemental Figure 10

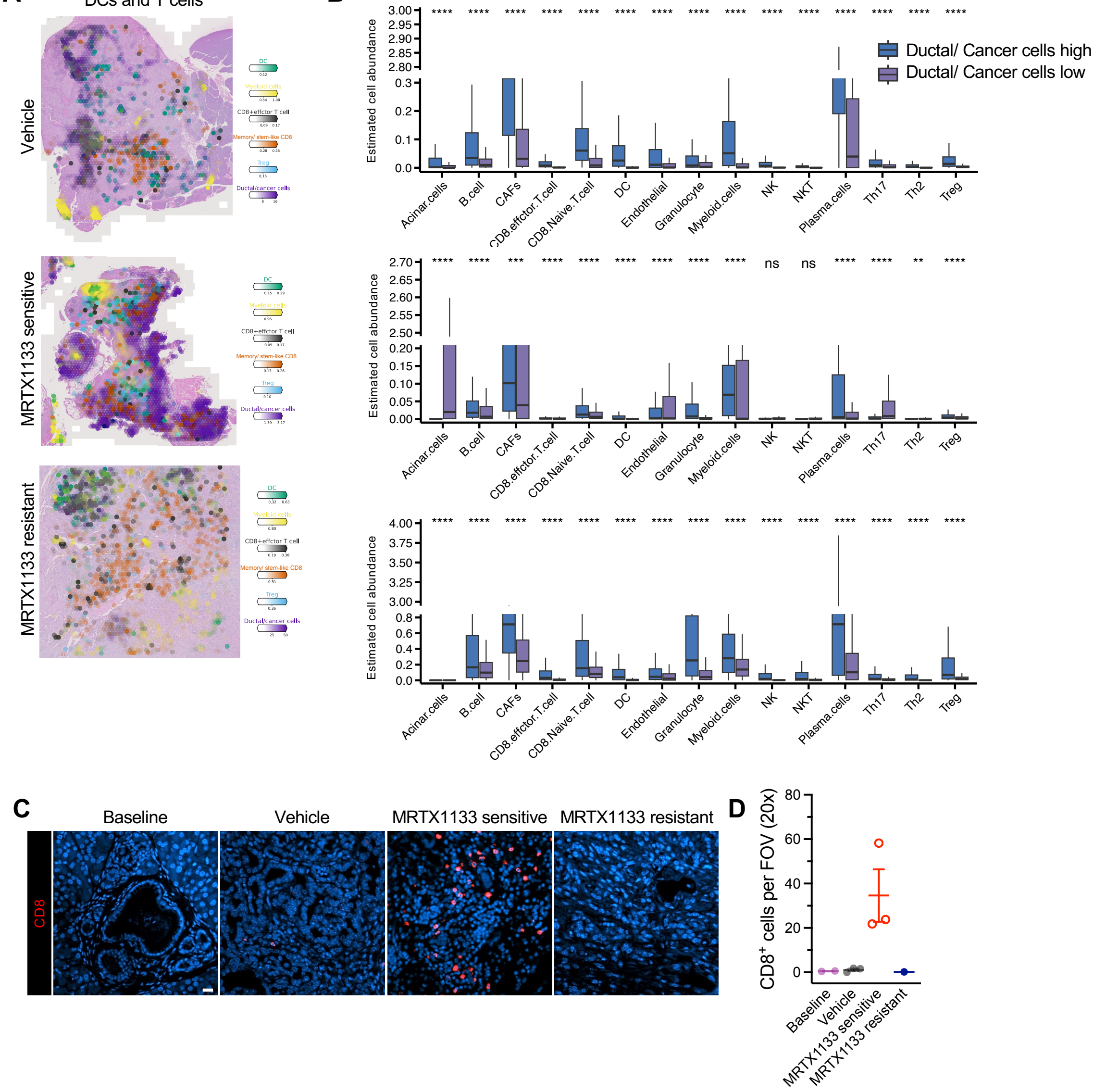


**Supplemental Figure 10: MRTX1133 resistance alters the tumor microenvironment.** (**A-B**) Spatial transcriptomics of orthotopic PKC-HY19636 tumors. Representative images (**A**) and quantification of cell abundance (**B**). Spots were stratified by ductal/cancer cell high and low areas. Vehicle, n=3; MRTX1133 sensitive, n=3; MRTX1133 resistant, n=3. (**C-D**) Representative images (**C**) and quantification (**D**) of CD8 immunostaining (red) of KPCY tumors. Baseline, n=2; vehicle, n=3; MRTX1133 sensitive, n=3; MRTX1133 resistant, n=1. Scale bar, 20 μm. Two-sided Mann-Whitney test was performed in (B). Data are presented as mean ± s.e.m. except for in (B) where it is presented as median and range. * $P < 0.05$, ** $P < 0.01$, **** $P < 0.0001$, *ns*: not significant.

#### Supplemental Figure 11

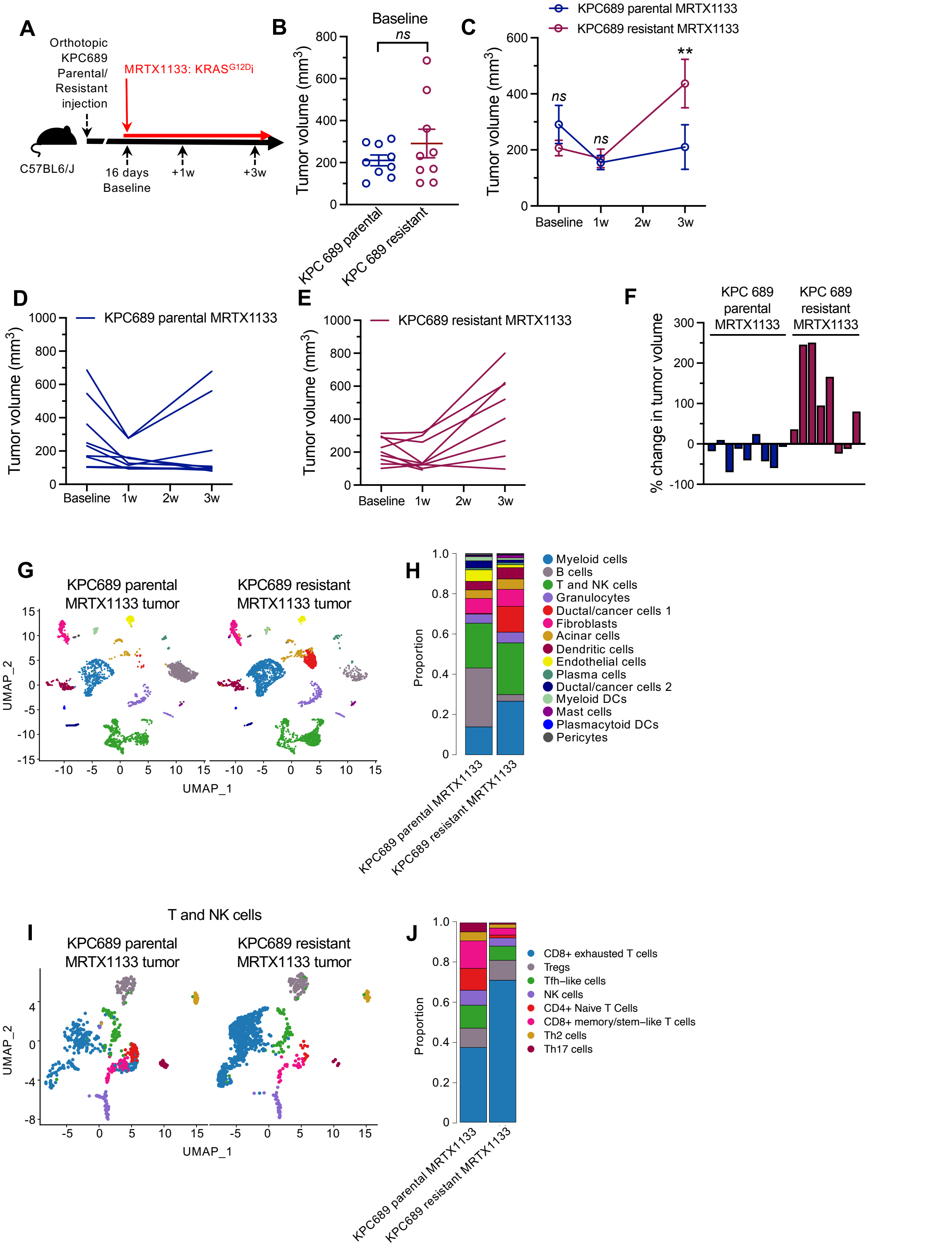


**Supplemental Figure 11: MRTX1133 resistant cells generated in culture lack**

**response to MRTX1133 in vivo and remodel the TME.** (**A**) Schematic of orthotopic KPC689 parental and KPC689 resistant tumors treated with MRTX1133. (**B-F**) Quantified MRI tumor volume at baseline (**B**) and following MRTX1133 treatment (**C**). Individual tumor volume curves for KPC689 parental (**D**), KPC689 resistant (**E**), and percent change in tumor volume at 3 weeks compared to baseline (**F**). KPC689 parental MRTX1133, n=9; KPC689 resistant MRTX1133, n=9. (**G-H**) UMAP (**G**) and relative proportions of all cells (**H**) of orthotopic KPC689 parental and resistant tumors treated with MRTX1133 determined by scRNA-seq**.** (**I-J**) UMAP (**I**) and relative proportions of T and NK cells (**J**) of orthotopic KPC689 parental and resistant tumors treated with MRTX1133 determined by scRNA-seq**.** Pancreata/tumors from 3 mice pooled per group. *ns*: not significant. Mixed-effects analysis with Bonferroni's multiple comparisons test performed in (C) and unpaired two-tailed t-test performed in (B). ** P <0.01, *ns*: not significant.

#### Supplemental Figure 12

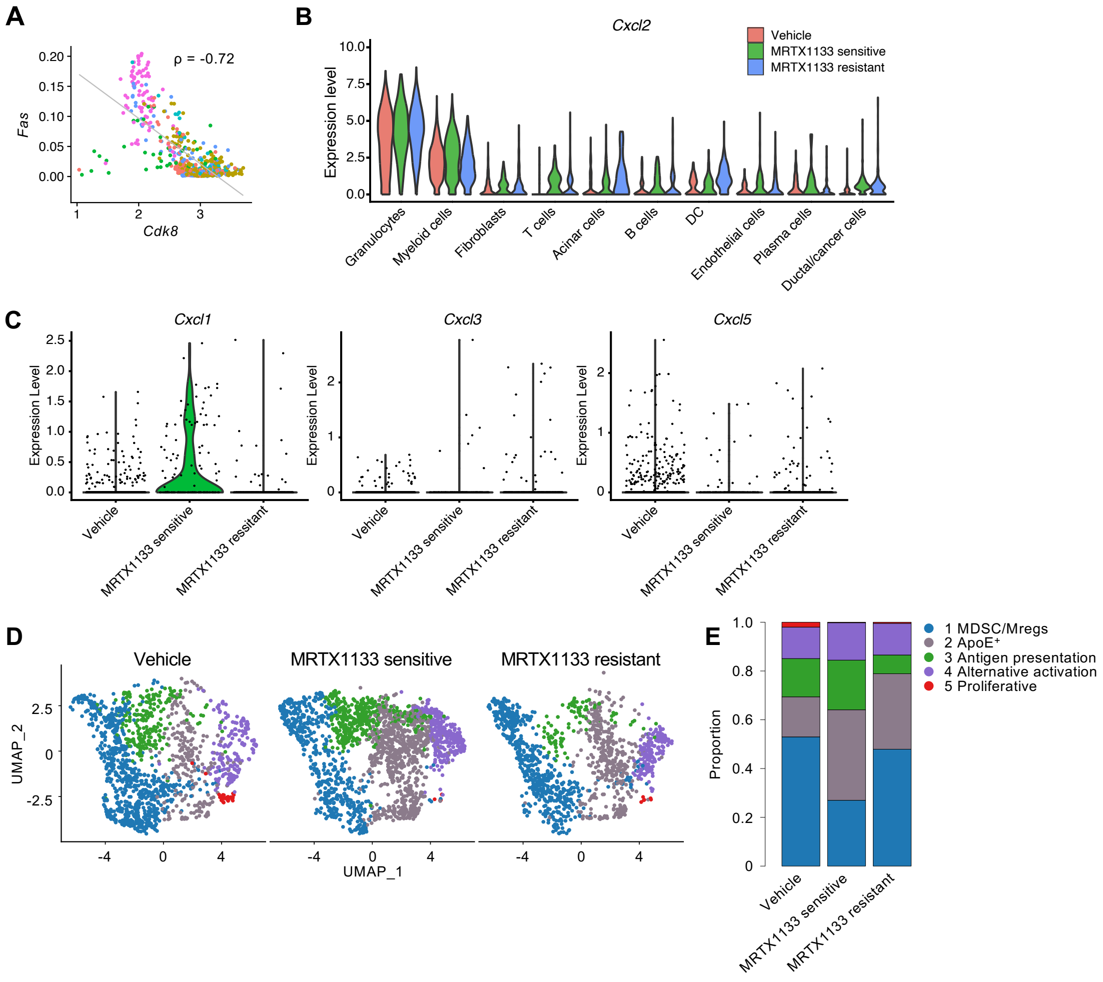


**Supplemental Figure 12: Myeloid infiltrate of MRTX1133 resistant tumors**. (**A**) Correlation between *Fas* and *Cdk8* expression based on scRNA-seq analysis. (**B**) *Cxcl2* expression in the expression of the indicated cell clusters of orthotopic PKC-HY19636 tumors determined by scRNA-seq. (**C**) Violin plots of *Cxcl1, Cxcl3, and Cxcl5* in ductal/cancer cells of orthotopic PKC-HY19636 tumors determined by scRNA-seq. (**D-E**) UMAP (**D**) and relative proportions of myeloid cells (**E**) of orthotopic PKC-HY19636 tumors determined by scRNA-seq. Data are reported as exact values except for in (B) where they are reported as violin plots.

#### Supplemental Figure 13

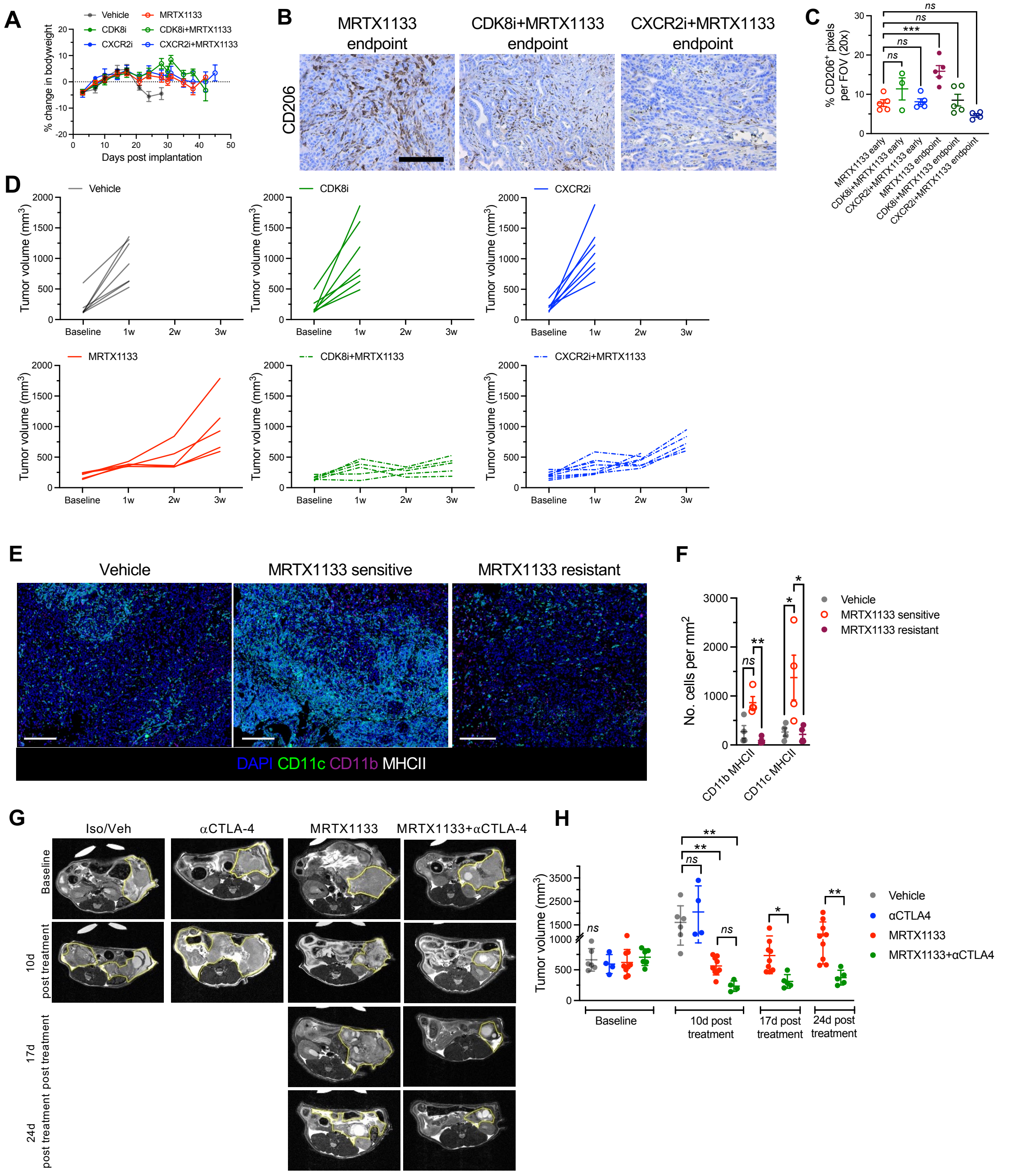

**Supplemental Figure 13: Analysis of immune infiltration and MRI analysis of PDAC mice.** (**A**) Bodyweight of mice bearing orthotopic PKC-HY19636 tumors post tumor-implantation. Vehicle, n=12; CDK8i, n=12; CXCR2i, n=12; MRTX1133, n=13; CDK8i+MRTX1133, n=12; CXCR2i+MRTX1133, n=13. (**B-C**) Representative CD206 staining (**B**) and quantification of orthotopic PKC-HY19636 tumors at endpoint (**C**). MRTX1133 early, n=5; CDK8i+MRTX1133 early, n=3; CXCR2i+MRTX1133 early, n=5; MRTX1133 endpoint, n=5; CDK8i+MRTX1133 endpoint, n=5; CXCR2i+MRTX1133 endpoint, n=4. Scale bar, 100 µm. Data from Fig. 5H are presented again here to directly compare early vs. endpoint tumors. (**D**) Tumor volumes measured by MRI of orthotopic PKC-HY19636 tumors treated with MRTX1133, CDK8i, and CXCR2i. Vehicle, n=7; CDK8i, n=7; CXCR2i, n=7; MRTX1133, n=6; CDK8i+MRTX1133, n=8; CXCR2i+MRTX1133, n=8. (**E-F**) Representative CODEX images of CD11c, CD11b, and MHCII (**E**) and quantification of cell abundance per tissue area of PKC-HY19636 orthotopic tumors (**F**). Vehicle, n=4; MRTX1133 sensitive, n=4; MRTX1133 resistant, n=4. (**G-H**) Representative images (**G**) and quantified tumor volume measured by MRI relative to tumor volume at 2 weeks following MRTX1133 treatment initiation (**H**). Yellow lines demarcate tumor regions. Veh, n=5; αCTLA-4, n=4; MRTX1133, n=9; MRTX1133+αCTLA-4, n=6. Kruskal-Wallis with Dunn's multiple comparisons test performed for (C), CD11b MHCII in (F), one-way ANOVA with Dunnett's multiple comparisons test performed for CD11c MHCII in (F) and 10d post-treatment comparisons in (H), and unpaired T-test for other comparisons in (H). Data are presented as mean ± s.e.m. except in (D) where individual mice are plotted and in (H) where mean ± s.d. is plotted. * $P<0.05$, ** $P<0.01$, *** $P<0.001$, *ns*: not significant.

Supplemental Figure 14

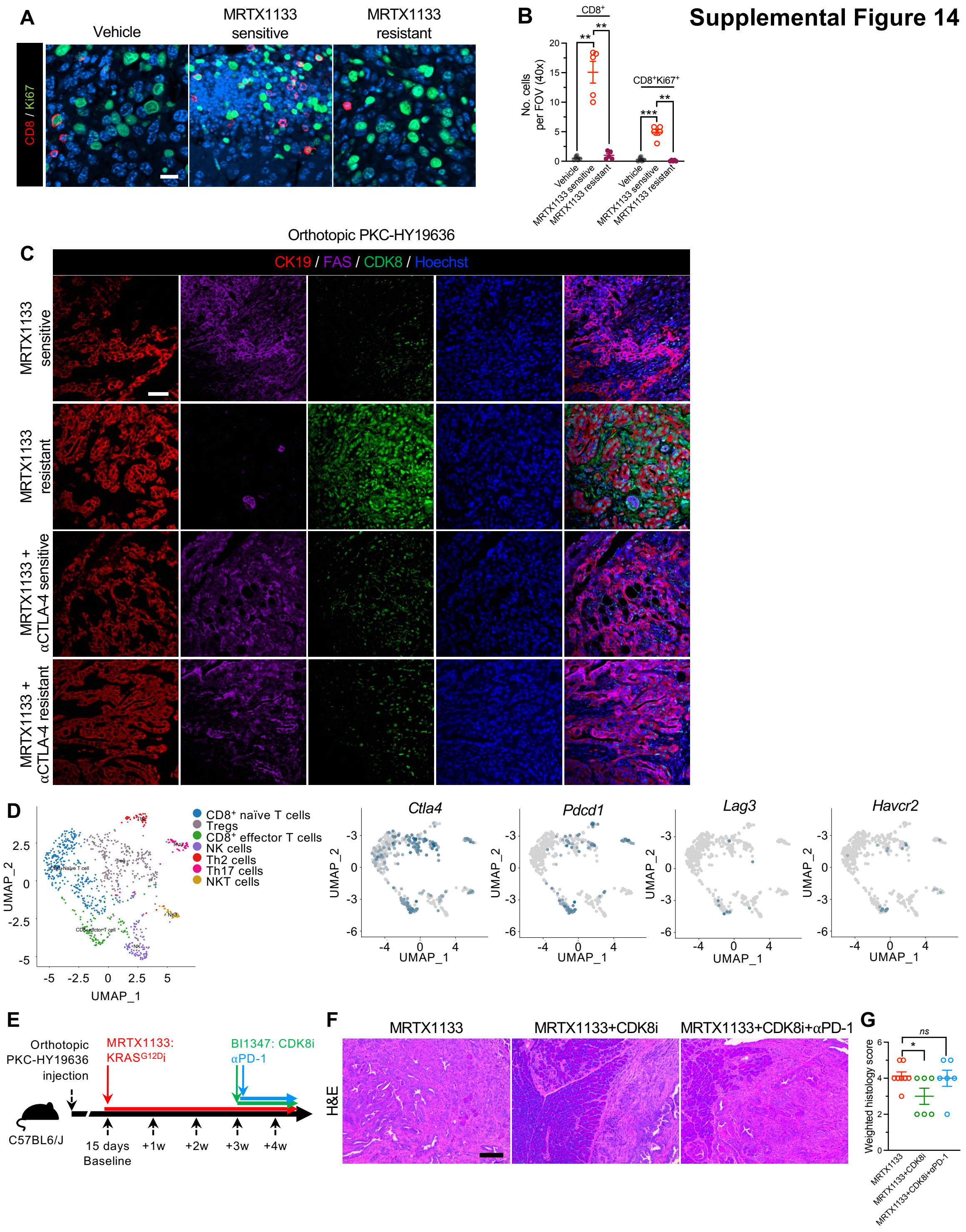


**Supplemental Figure 14: T cell responses in resistant tumors.** (**A-B**) Representative images (**A**) and quantification of $CD8^+$ and $CD8^+Ki67^+$ cells (**B**). Vehicle, n=5, MRTX1133

sensitive, n=5, MRTX1133 resistant, n=5. Scale bar, 20 μm. (**C**) Single channel representative images of CK19 (red), FAS (purple), and CDK8 (green) stained orthotopic PKC-HY19636 tumors. Images from Fig. 5F are presented again here with each channel included individually. Scale bar, 50 μm. (**D**) UMAP of T cells of orthotopic PKC-HY19636 tumors and expression of checkpoint molecules determined by scRNA-seq. (**E**) Schematic of orthotopic PKC-HY19636 tumors treated with MRTX1133, CDK8i (BI1347), and αPD-1. (**F-G**) Representative H&E images (**F**) and quantification (**G**) of orthotopic PKC-HY19636 tumors. MRTX1133, n=8; MRTX1133+CDK8i, n=6; MRTX1133+CDK8i+αPD-1, n=6. Scale bar, 100 μm. Brown-Forsythe and Welch ANOVA with Dunnett T3's multiple comparisons test performed in (B). Chi-square test performed comparing responsive (score≤2) to non-responsive (score>2) in (G). Data are presented as mean ± s.e.m. * P< 0.05, ** P < 0.01, *ns*: not significant.

#### Supplemental Figure 15

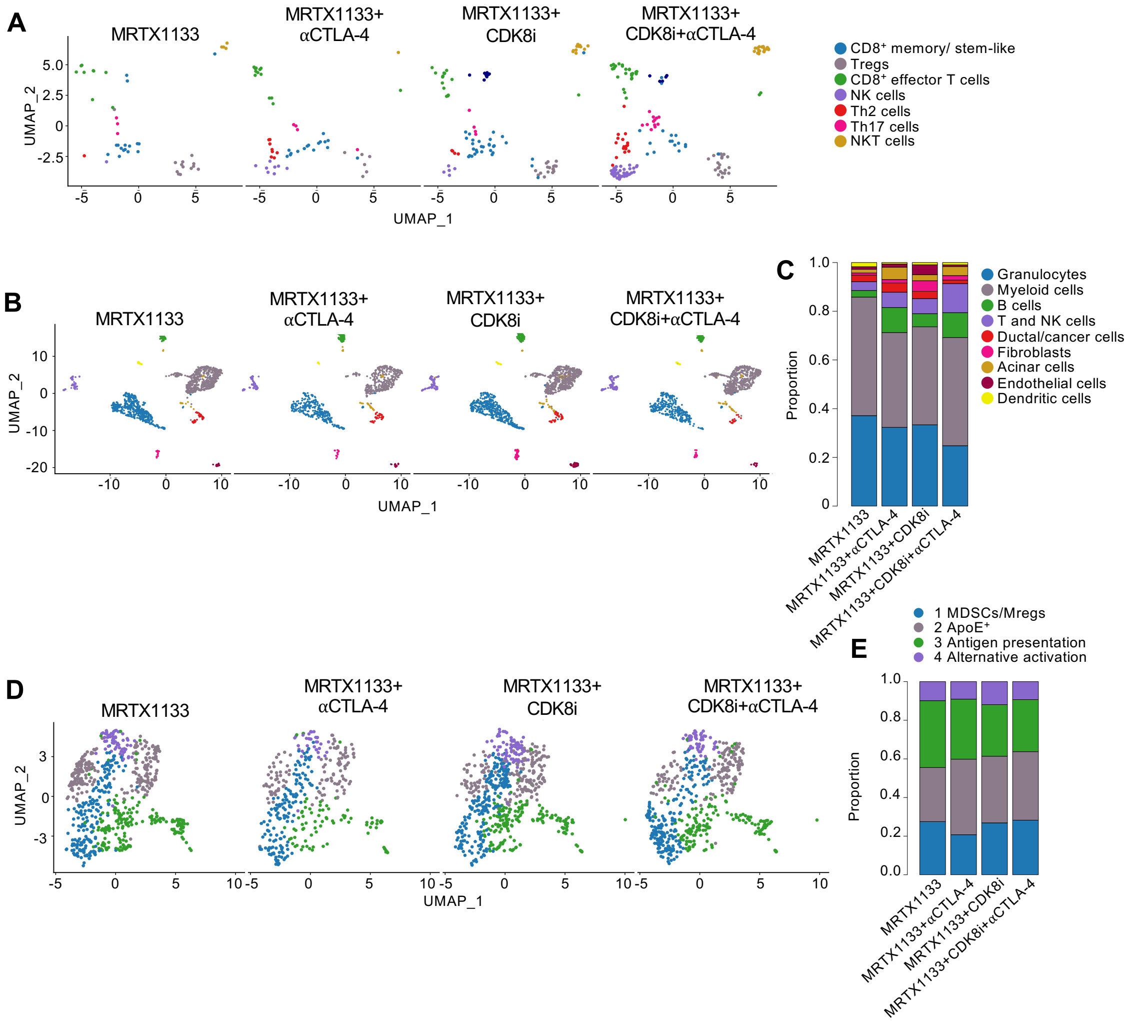


**Supplemental Figure 15: T cell and myeloid cell infiltration in MRTX1133 resistant tumors treated with CDK8i with αCTLA-4**. (**A**) UMAP of T cell subsets of PKC-HY19636 tumors analyzed by scRNA-seq. (**B-C**) UMAP (**B**) and relative proportions of cells (**C**) of orthotopic PKC-HY19636 tumors treated with MRTX1133, CDK8i, and αCTLA-4 determined by scRNA-seq. (**D-E**) UMAP (**D**) and relative proportions (**E**) of myeloid subsets in orthotopic PKC-HY19636 tumors analyzed by scRNA-seq. Data are reported as exact values (C) and (E).

#### Supplemental Tables

**Supplemental Table 1: CNVs in orthotopic PKC-HY19636 tumors and MRTX1133 resistant cell lines**. (**A**) CNVs detected in orthotopic PKC-HY19636 tumors. (**B**) CNVs detected in MRTX1133 in vitro resistant KPC689 and HY19636 cell lines expressed as a relative ratio to parental cell lines. (**C**) CNVs detected in MRTX1133 in vitro resistant KPC689 and HY19636 cell lines expressed as original copy numbers.

### Uncropped blots

##### Supplemental Figure 6A

Uncropped blots

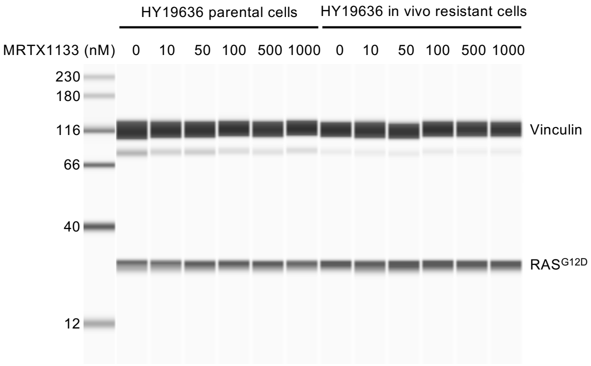


##### Supplemental Figure 6B

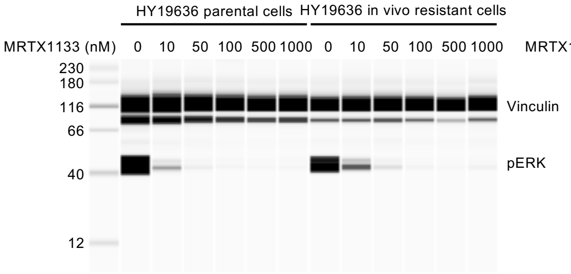


High contrast

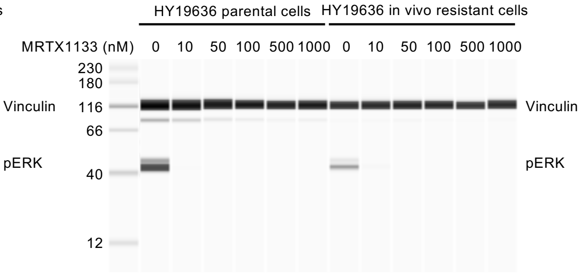


Low contrast

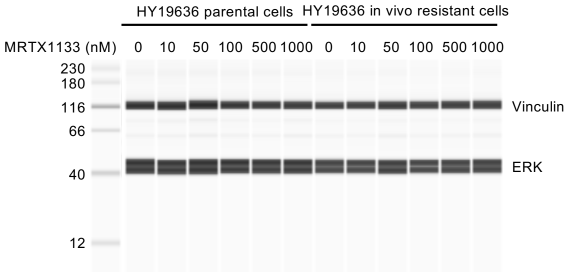

**Supplemental Figure 7C**

#### Uncropped blots

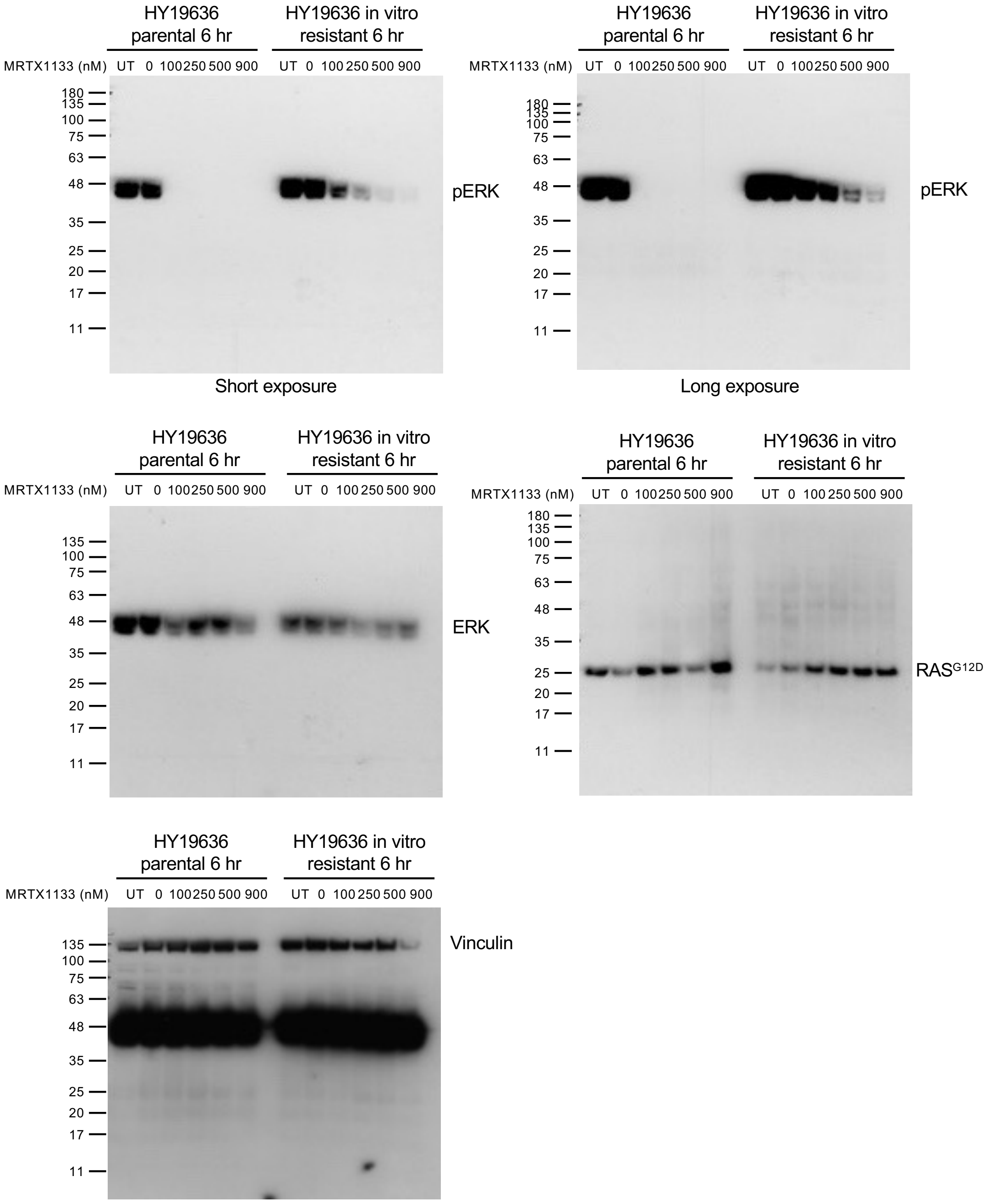

**Supplemental Figure 7C**

#### Uncropped blots

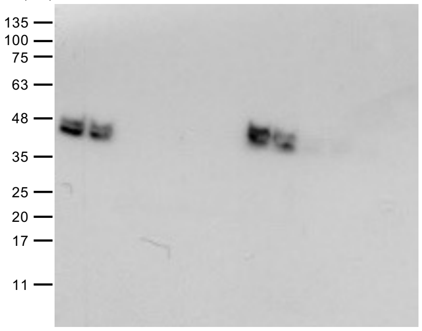


Short exposure

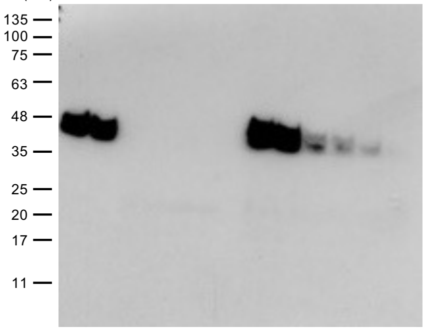


Long exposure

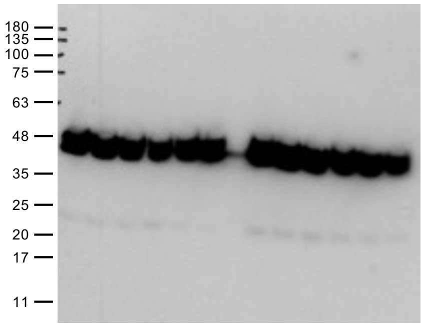


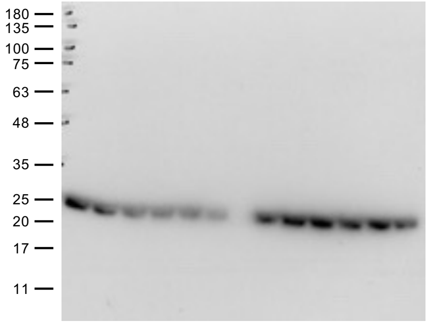


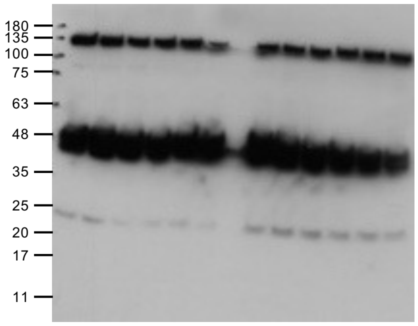

**Supplemental Figure 7C**

#### Uncropped blots

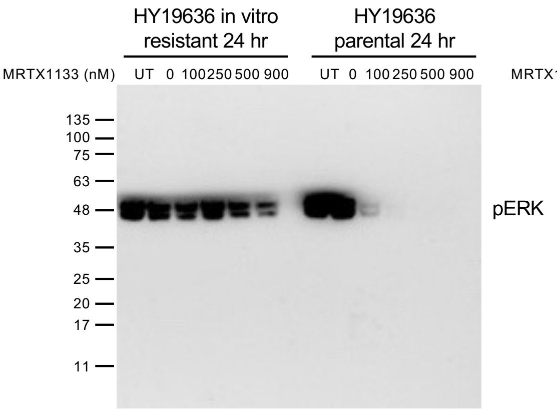

Short exposure

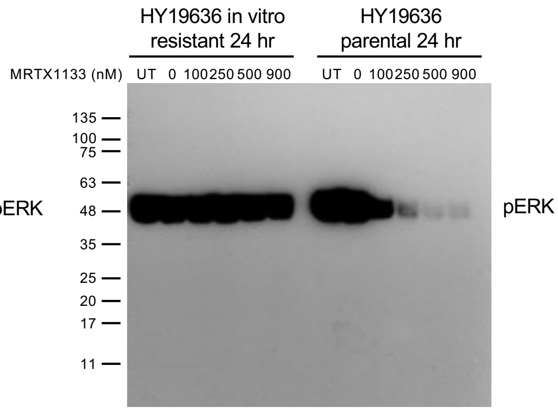

Long exposure

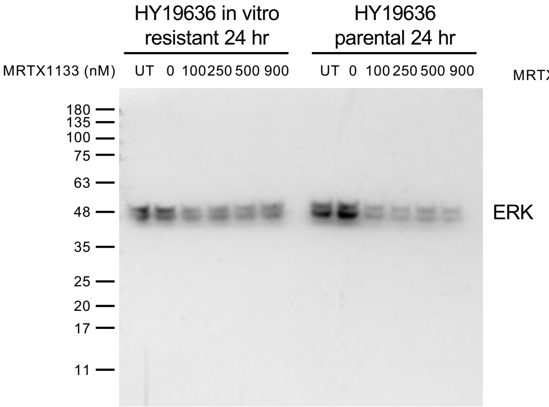


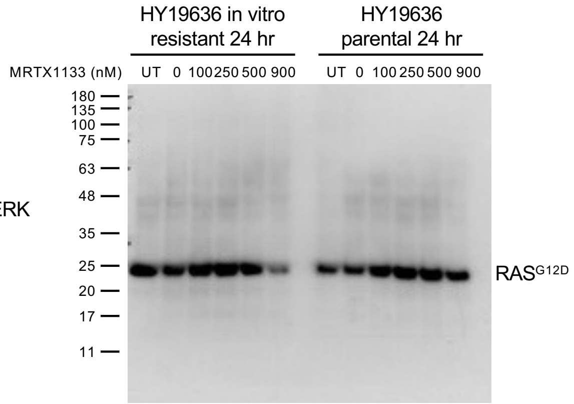


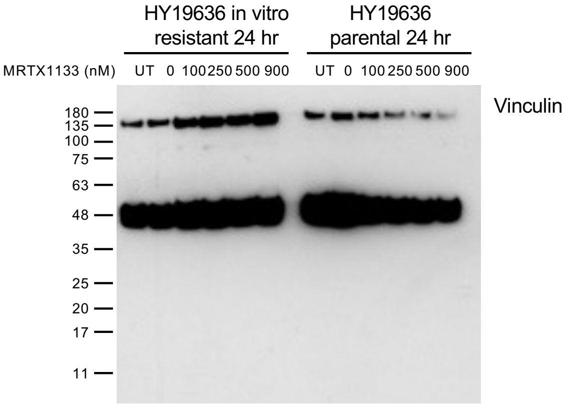

**Supplemental Figure 7E**

#### Uncropped blots

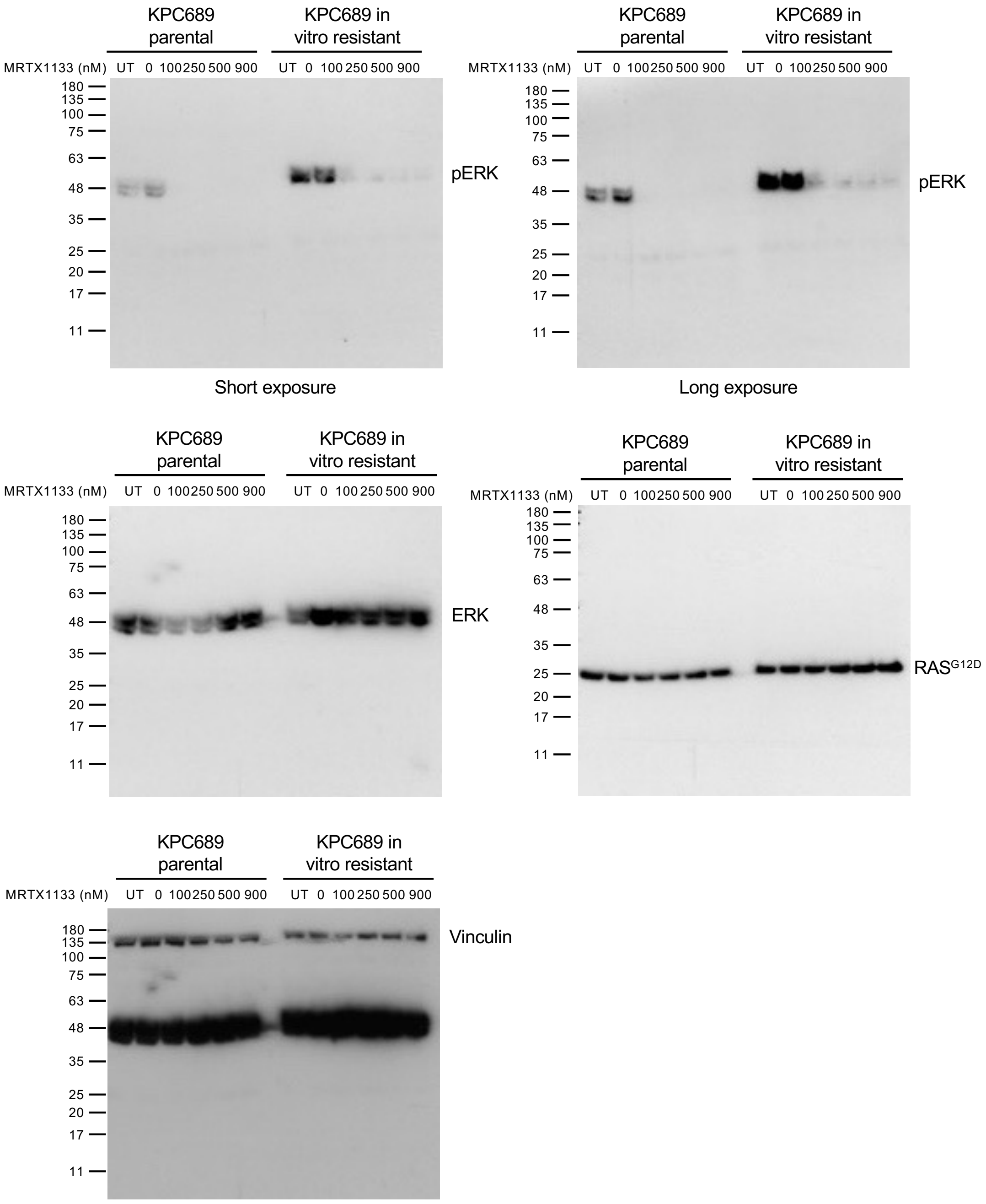
